# Intersection of phosphate transport, oxidative stress and TOR signalling in *Candida albicans* virulence: Pho84 in *Candida albicans* virulence

**DOI:** 10.1101/317933

**Authors:** Ning-Ning Liu, Priya Uppuluri, Achille Broggi, Angelique Besold, Kicki Ryman, Hiroto Kambara, Norma Solis, Viola Lorenz, Wanjun Qi, Maikel Acosta-Zaldivar, S. Noushin Emami, Bin Bao, Dingding An, Francisco A. Bonilla, Martha Sola-Visner, Scott G. Filler, Hongbo R. Luo, Ylva Engström, Per Olof Ljungdahl, Valeria C. Culotta, Ivan Zanoni, Jose L. Lopez-Ribot, Julia R. Köhler

**Affiliations:** Department of Infectious Diseases, Boston Children’s Hospital/Harvard Medical School, Boston, MA 02115, USA; School of Public Health, Shanghai Jiao Tong University School of Medicine, Shanghai 200025, China; Division of Infectious Diseases, Los Angeles Biomedical Research Institute at Harbor-UCLA Medical Center, Torrance, CA 90502, USA; Division of Gastroenterology, Boston Children’s Hospital/Harvard Medical School, Boston, MA 02115, USA; Department of Biochemistry and Molecular Biology, The Johns Hopkins University Bloomberg School of Public Health, Baltimore, MD 21205, USA; The Wenner-Gren Institute, Stockholm University, Stockholm SE-106 91, Sweden; Department of Pathology, Joint Program in Transfusion Medicine, Boston Children’s Hospital/Harvard Medical School, Boston, MA 02115, USA; Department of Newborn Medicine, Boston Children’s Hospital/Harvard Medical School, Boston, MA 02115, USA; Clinical Immunology Program, Boston Children’s Hospital, Boston, Massachusetts 02115, USA; Department of Biotechnology and Biosciences, University of Milano-Bicocca, Piazza della Scienza 2, 20126 Milan, Italy; Department of Biology, University of Texas at San Antonio, San Antonio, Texas 78249, USA

## Abstract

Phosphate is an essential macronutrient required for cell growth and division. Pho84 is the major high-affinity cell-surface phosphate importer of *Saccharomyces cerevisiae* and a crucial element in the phosphate homeostatic system of this model yeast. We found that loss of *Candida albicans* Pho84 attenuated virulence in *Drosophila* and murine oropharyngeal and disseminated models of invasive infection, and conferred hypersensitivity to neutrophil killing. Susceptibility of cells lacking Pho84 to neutrophil attack depended on reactive oxygen species (ROS): *pho84-/-* cells were no more susceptible than wild type *C. albicans* to neutrophils from a patient with chronic granulomatous disease, or to those whose oxidative burst was pharmacologically inhibited or neutralized. *pho84-/-* mutants hyperactivated oxidative stress signalling. They accumulated intracellular ROS in the absence of extrinsic oxidative stress, in high as well as low ambient phosphate conditions. ROS accumulation correlated with diminished levels of the unique superoxide dismutase Sod3 in *pho84-/-* cells, while *SOD3* overexpression from a conditional promoter substantially restored these cells’ oxidative stress resistance in vitro. Repression of *SOD3* expression sharply increased their oxidative stress hypersensitivity. Neither of these oxidative stress management effects of manipulating *SOD3* transcription was observed in *PHO84* wild type cells. Sod3 levels were not the only factor driving oxidative stress effects on *pho84-/-* cells, though, because overexpressing *SOD3* did not ameliorate these cells’ hypersensitivity to neutrophil killing ex vivo, indicating Pho84 has further roles in oxidative stress resistance and virulence. Measurement of cellular metal concentrations demonstrated that diminished Sod3 expression was not due to decreased import of its metal cofactor manganese, as predicted from the function of *S. cerevisiae* Pho84 as a low-affinity manganese transporter. Instead of a role of Pho84 in metal transport, we found its role in TORC1 activation to impact oxidative stress management: overexpression of the TORC1-activating GTPase Gtr1 relieved the Sod3 deficit and ROS excess in *pho84-/-* null mutant cells, though it did not suppress their hypersensitivity to neutrophil killing or hyphal growth defect. Pharmacologic inhibition of Pho84 by small molecules including the FDA-approved drug foscarnet also induced ROS accumulation. Inhibiting Pho84 could hence support host defenses by sensitizing *C. albicans* to oxidative stress.

## Author Summary

*Candida albicans* is the species most often isolated from patients with invasive fungal disease, and is also a common colonizer of healthy people. It is well equipped to compete for nutrients with bacteria co-inhabiting human gastrointestinal mucous membranes, since it possesses multiple transporters to internalize important nutrients like sugars, nitrogen sources, and phosphate. During infection, the fungus needs to withstand human defense cells that attack it with noxious chemicals, among which reactive oxygen species (ROS) are critical. We found that a high-affinity phosphate transporter, Pho84, is required for *C. albicans’* ability to successfully invade animal hosts and to eliminate ROS. Levels of a fungal enzyme that breaks down ROS, Sod3, were decreased in cells lacking Pho84. A connection between this phosphate transporter and the ROS-detoxifying enzyme was identified in the Target of Rapamycin (TOR) pathway, to which Pho84 is known to provide activating signals when phosphate is abundant. Small molecules that block Pho84 activity impair the ability of *C. albicans* to detoxify ROS. Since humans manage phosphate differently than fungi and have no Pho84 homolog, a drug that inhibits Pho84 could disable the defense of the fungus against the host.

## Introduction

*Candida albicans* is the most common invasive human fungal pathogen, whose infections carry a high mortality rate (1). It is also a widespread commensal, colonizing gastrointestinal mucous membranes of around half of healthy humans (2) and competing with myriad bacteria for nutrients shed by the host or extractable from the food stream (3, 4). Sources of the macronutrients carbon, nitrogen and phosphate must be distributed between the host and its bacterial and fungal colonizers. During invasive disease, *C. albicans* uses the human as its source of nutrients and must withstand the host immune system (1).

Availability of inorganic phosphate (Pi) is critical for cells metabolizing carbon and nitrogen sources, synthesizing ribosomes and membranes, and preparing for DNA replication. Bacteria devote a Pi signalling and acquisition system, the PHO regulon, to Pi homeostasis. In many pathogenic bacteria, the PHO regulon has been linked to virulence, though definition of the perturbed pathogenic mechanisms has often remained elusive (5). In some bacteria like *Vibrio cholerae*, the PHO regulon’s transcriptional regulator also controls or co-regulates hundreds of genes related to virulence and unrelated to Pi homeostasis (6, 7) so that the role of the PHO regulon may be to inform the bacterial cell of the presence of host signals in its environment. In fungi and plants, a more complex PHO regulon monitors the availability of intracellular as well as extracellular Pi and is integrated with a large signalling network responding to nutrient availability and the absence of stressors (8).

Unicellular parasitic human pathogens, evolved to invade the bloodstream and infect vital tissues, have Pi import systems whose importance has recently come into focus (9). Significantly, *Leishmania infantum*, an intracellular parasitic agent of human visceral leishmaniasis which is a neglected tropical disease of 12 million people (10), expresses a Pi transporter homologous to the *Saccharomyces cerevisiae* high-affinity H+-Pi symporter Pho84 as its major mode of Pi acquisition (11). Insect stages of the related kinetoplastid parasite *Trypanosoma cruzi*, the agent of Chagas disease estimated to chronically infect 8 million people while leading to inexorable cardiac death in 20-30% of the infected (12, 13), are Pi-dependent for their cellular differentiation (14). *T. cruzi* insect stages utilize homologs of the *Saccharomyces cerevisiae* high- and low-affinity Pi importers to acquire the Pi quantities that permit their development and proliferation (14).

A high-affinity Pi transporter of *C. albicans*, Pho84, is required for normal Target of Rapamycin (TOR) signalling and hyphal morphogenesis (8). Pharmacologic inhibition of Pho84 potentiates the activity of two major antifungal classes (8). Pho84 is one of four predicted *C. albicans* plasma membrane Pi transporters, so that redundancy of its activity would be expected, and a role of Pho84 in virulence could not be assumed a priori. We previously observed failure of *pho84* deletion mutants to appropriately induce hyphal growth in response to several in vitro conditions (8). Since hyphal growth is a known virulence determinant in *C. albicans*, we examined virulence of *pho84* mutant cells first in a wild type *Drosophila melanogaster* model, then in two murine models. Despite its redundancy as a Pi transporter, mutants in Pho84 exhibited attenuated virulence in these models, which may partially be attributable to their hyphal morphogenesis defect, observable in one of the murine models. Finding a requirement for Pho84 in resistance of *C. albicans* cells to whole human blood exposure, we then focused on isolating a molecular mechanism of this role of Pho84. Pho84 mutants were hypersensitive to killing by human neutrophils, the major cellular blood component that controls invading fungi. Our analysis led to the complex *C. albicans* superoxide dismutase (SOD) system (15), and demonstrated a novel requirement for active TOR complex 1 (TORC1) in maintaining appropriate cytosolic SOD expression. Generation of the superoxide anion and other reactive oxygen species (ROS) is a major defense mechanism of host phagocytes against the fungus (16). In addition to its role in hyphal growth, the link between a cytosolic SOD and TORC1 activity provides a rationale for the contribution of Pho84, a redundant member of the cell surface Pi importers, to virulence of *C. albicans.*

## Results

### Pho84 is required for virulence of *C. albicans* in a wild-type *Drosophila* model

Null mutants in *PHO84* are defective in TORC1 signaling and in hyphal growth (8). As *C. albicans* hyphal growth is a virulence determinant, we examined *pho84* null mutant (-/-, *pho84/pho84)* cells’ virulence in a *Drosophila melanogaster* model (17). In the wild type OregonR strain we used, *Drosophila* immune responses are intact, so that the fungus confronts the full complement of innate immunity. By 5 days after infection, 30% of flies injected with *PHO84* wild type, and 8% of those injected with *pho84* null mutant cells had died (*p*<0.001 by Kaplan-Meier analysis). Virulence behavior of reintegrant (-/-/+, *pho84/pho84::PHO84*) cells was statistically indistinguishable from wild type (*p*=0.3) (Fig. 1A). Hence, in a model of simple innate immunity, Pho84 was required for virulence.

**Figure 1.**
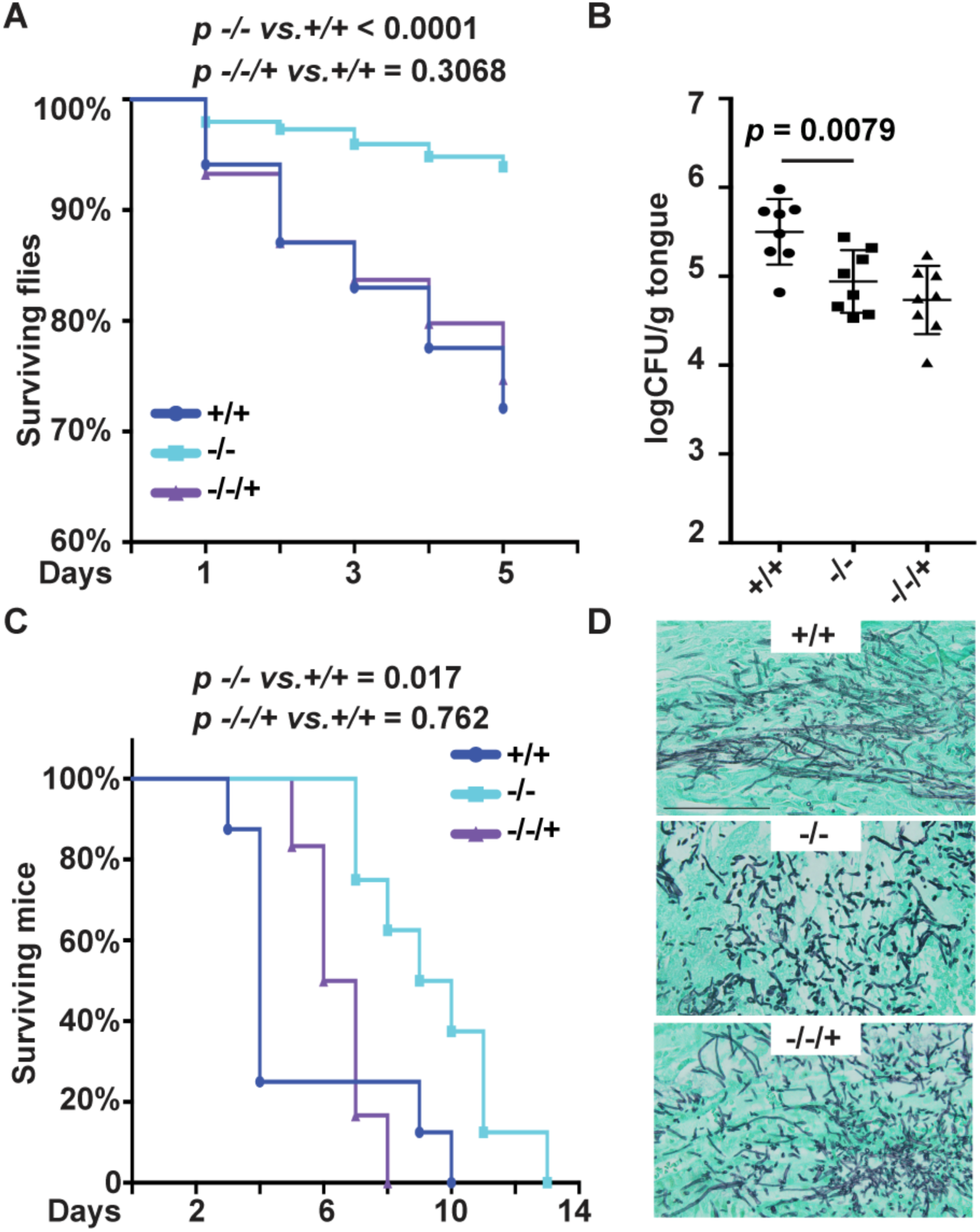
Virulence is attenuated in *C. albicans* cells lacking *PHO84.* (A) Percentage of surviving *Drosophila melanogaster* after infection with cells of the *PHO84* genotypes +/+, JKC915; -/-, JKC1450 and -/-/+, JKC1588 respectively. The flies were injected with 50 nl fungal cell suspensions with a micro-injector. At least 3 (up to 7) biological replicates and 6 -15 technical replicates per strain were performed. (B) Oral fungal burden of mice with oropharyngeal candidiasis was calculated by enumerating CFU per gram tongue tissue after 5 days of infection. Each dot represents the tissue fungal burden of one mouse. (C) Kaplan-Meier survival plot of mice with disseminated candidiasis. Eight mice per strain were injected with strains as in A respectively. (D) Kidneys from *+/+, -/-* and -/-/+ infected mice were isolated and sections stained with Gomori Methenamine Silver. Size bar 0.1 mm.

### Pho84 contributes to virulence in murine disseminated and oropharyngeal candidiasis

Given their attenuation in an insect model, we examined the virulence of *pho84* null mutant cells in two murine models of infection. One path of natural *C. albicans* infection is invasion of the mucosa. We compared *pho84* null and wild type cells for the ability to proliferate in a murine oropharyngeal candidiasis model (18). Loss of Pho84 resulted in a decreased fungal burden of tongue tissue (Fig. 1B), indicating that in this model, functions of Pho84 contribute to virulence during oropharyngeal infection.

Loss of virulence of the *pho84* null and *pho84-/-/+* reintegrant cells in this model was similar. We previously observed haploinsufficiency of *PHO84* heterozygous and reintegrant cells (8). In fact, in one phenotype, hypersensitivity to oxidative stress, the degree of haploinsufficiency correlated with the intensity of the stress (S1 Fig.). Nevertheless, we cannot exclude that a second mutation in the null mutant strain diminished its virulence in this model.

Once *C. albicans* cells have crossed the mucosa and entered the bloodstream, they disseminate to distant organs and initiate new foci of infection. We asked whether Pho84 is required for *C. albicans* virulence during hematogenous infection. Survival of mice injected intravenously with wild type, *pho84* null mutant and *pho84-/-/+* reintegrant cells was compared, and more than 70% of the mice infected with the wild type strain died within 4 days of infection (Fig. 1C). Loss of Pho84 significantly extended survival in this model; *p*=0.017 for wild type versus *pho84* null mutant cells, and *p*=0.76 for wild type versus reintegrant by Kaplan-Meier analysis. We assessed the morphogenetic state of infecting filamentous cells according to the criteria defined by Sudbery et al. (19). Gomori-Methenamine Silver stained sections of kidneys from moribund mice showed wild type cells growing predominantly in the hyphal form with interspersed yeast cells (Fig. 1D). In contrast, *pho84* null mutant cells were a mixture of pseudohyphal filaments and yeast with fewer hyphae (Fig. 1D), reflecting defective hyphal growth previously seen in vitro (8), while reintegrant cells grew as hyphae, pseudohyphae and yeast (Fig. 1D). Hence, in a model of disseminated disease, loss of Pho84 correlated with prolonged survival of infected animals, and a hyphal growth defect of null mutant cells may have contributed to their virulence defect.

### Pho84 is required for resistance to whole blood candidacidal activity, neutrophil killing and reactive oxygen species (ROS) exposure

During bloodstream infection, invading *C. albicans* cells encounter host blood components. We incubated heparinized whole blood from healthy human volunteers with *C. albicans* and found survival of *pho84* null mutant cells to be significantly decreased after 5 hours, compared with the wild type and reintegrant (Fig. 2A). This finding indicates that Pho84 has a role in *C. albicans’* tolerance of whole blood candidacidal activity.

**Figure 2.**
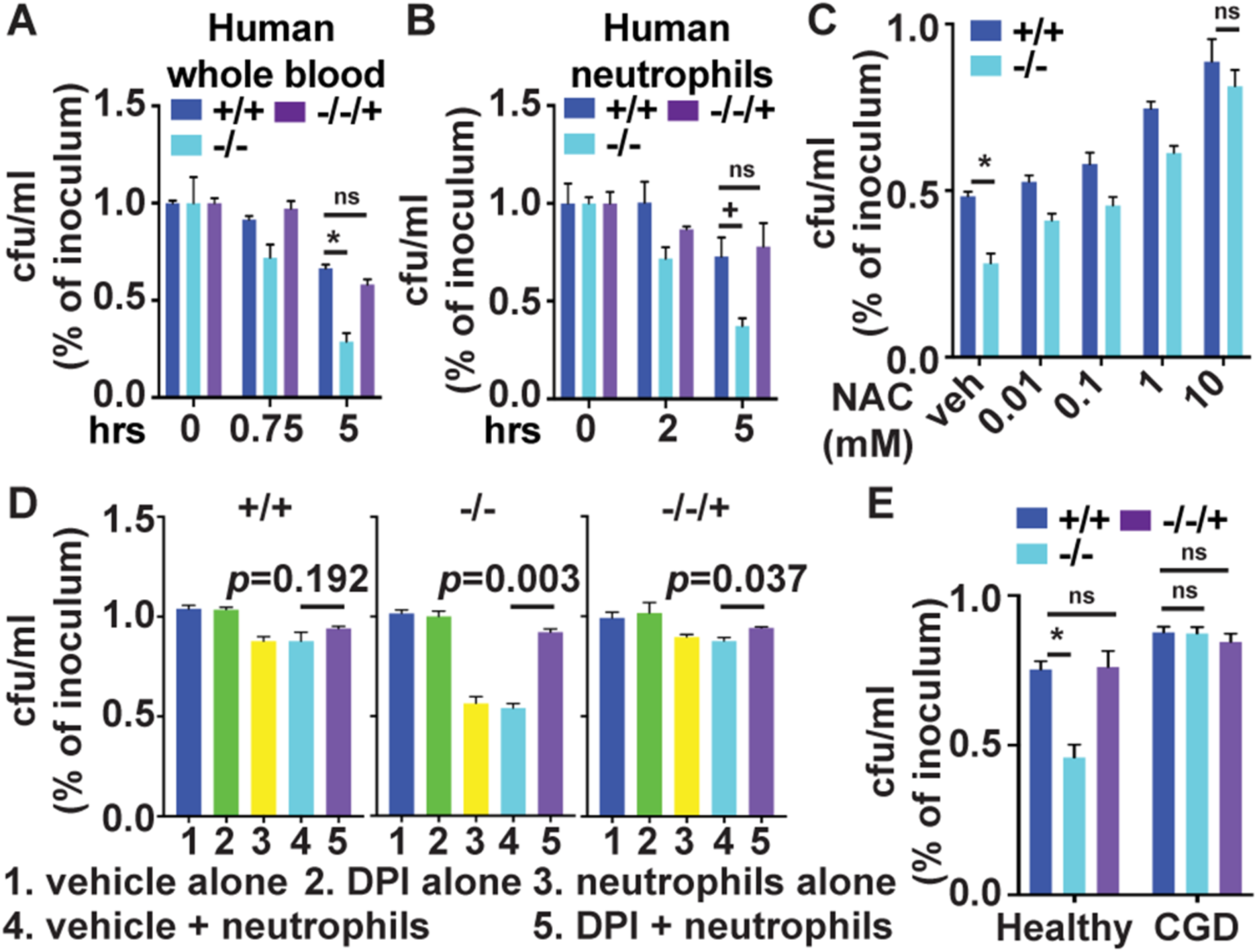
Pho84 is required for resistance to killing by whole blood or neutrophils, in dependence on neutrophil ROS. (A) Percent survival of *C. albicans* cells after incubation with whole blood from healthy human volunteers; cells of genotypes +/+, JKC915; -/-, JKC1450 and -/-/+, JKC1588 were inoculated into blood and plated onto agar medium at the indicated time points for calculation of CFU/ml. *-/-* versus *+/+* at 5 h *p*=0.008. (B) Percent survival of *C. albicans* cells after incubation with neutrophils, strains as in *A*, at M.O.I. 2 for 2 hrs and 5 hrs. *-/-* versus *+/+* at 5 h *p*=0.04. (C) Human peripheral blood-derived neutrophils pretreated with different concentrations of N-acetyl-l-cysteine (NAC) were incubated for 90 min with strains as in A at M.O.I. 2. *-/-* versus *+/+* with vehicle *p*<0.0001. (D) Human peripheral blood-derived neutrophils pretreated with 10 μM Diphenyleneiodonium (DPI) were incubated for 90 min with strains as in A at M.O.I. 2. Vehicle alone, DPI alone and neutrophil alone groups are controls. (E) Chronic Granulomatous Disease patient-derived neutrophils (CGD) were incubated for 2 hours with strains as in A at M.O.I. 2. *p*<0.0001 for *-/-* in healthy control neutrophils (control) versus *+/+* in control, all others non-significant. *p* values per Student’s t-test. \**p*<0.01; +*p*<0.05; ns is non-significant. A-D representative of at least 3 biological replicates.

Neutrophils are the major contributors to *C. albicans’* transcriptional responses to whole blood exposure, and are critical components of cellular innate immunity against invasive candidiasis (20-22). We therefore tested the ability of *pho84* null mutant cells to survive the attack of human neutrophils, using the HL-60 human promyelocytic leukemia cell line which can differentiate into neutrophil-like cells (23). Null mutants in *PHO84* were significantly more sensitive to killing by these phagocytic cells than wild type or reintegrant cells (S2A Fig.). We confirmed this finding in primary neutrophils isolated from healthy human donors (Fig. 2B). To distinguish hypersensitivity of *pho84* null mutant cells to neutrophil cidal activity from their putatively increased phagocytic uptake, we performed phagocytosis assays and found that *pho84* null and wild type cells were equally taken up by the neutrophils (S2B Fig.). Similarly, we examined whether *pho84* null mutants may stimulate neutrophil ROS production more effectively, since ROS are a major neutrophil candidacidal mechanism (24). We found no difference in intracellular or extracellular ROS production between neutrophils interacting with wild type or reintegrant cells, and those interacting with *pho84* null mutant cells (S2C and S2D Figs). These findings, obtained with neutrophils from identified healthy volunteers and from random, unidentifiable blood bank donors, indicate that Pho84 contributes to protection of *C. albicans* from neutrophil killing.

To examine whether in fact ROS are responsible for *pho84* null mutant cells’ increased susceptibility to neutrophils’ cidal activity, we treated Candida-ingesting neutrophils with the ROS-scavenging compound N-acetyl cysteine (NAC) (25). In a dose-dependent manner, NAC rescued hypersensitivity of *pho84* null mutant cells to neutrophil killing (Fig. 2C). To block ROS production a priori, we then inhibited neutrophil NADPH oxidase (NOX), the enzyme complex responsible for generating the ROS oxidative burst, by preincubating neutrophils with diphenyliodonium (DPI) (26, 27) and followed survival of *C. albicans* cells. Inhibition of neutrophil NOX with DPI abolished hypersensitivity of *pho84* null mutant cells to neutrophil killing (Fig. 2D). Chronic granulomatous disease (CGD) is caused by mutations that disrupt NOX function. *pho84* null mutant cells were equally resistant to killing by neutrophils isolated from a CGD patient as wild type cells, while in the same experiment, as previously, they were hypersensitive to killing by neutrophils from a control healthy volunteer (Fig. 2E). These results suggest that Pho84 is required for resistance specifically to ROS-mediated neutrophil candidacidal activity (24).

These findings raised the question whether *pho84* null mutants are simply hypersensitive to ROS. Exposing wild type, *pho84* null, and reintegrant cells to inducers of superoxide anion, plumbagin and menadione, as well as to hydrogen peroxide (H_2_O_2_), we found that each of these compounds inhibited growth of the mutant more strongly than that of the wild type (Fig. 3A). Hypersensitivity of *pho84* null mutant cells to neutrophil killing may therefore be due to their hypersensitivity to oxidative stress.

**Figure 3.**
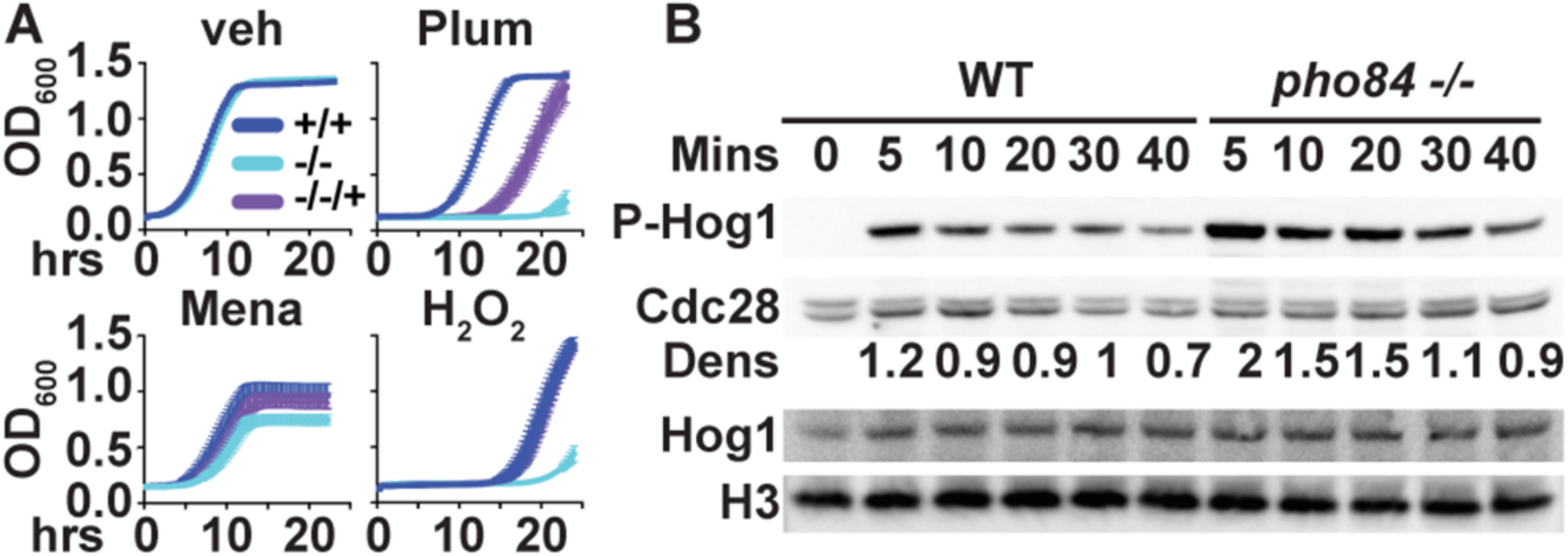
Resistance to oxidative stressors is decreased, and Hog1 phosphorylation is increased in cells lacking *PHO84.* (A) Cells from overnight liquid cultures were inoculated into YPD with vehicle (veh), 25 μM plumbagin (Plum), 40 μM menadione (Mena), and 40 mM H_2_O_2_ to OD_600_ 0.1. OD_600_ was monitored every 15 minutes for strains +/+, JKC915; -/-, JKC1450 and -/-/+, JKC1588. (B) Western blot of cells incubated in YPD containing 5mM H_2_O_2_ for 5 min, 10 min, 20 min, 30 min and 40 min. The membrane was probed for P-Hog1, total Hog1, and loading controls Cdc28 (anti PSTAIRE motif) and Histone H3. Dens: ratio of densitometry of the phosphorylated Hog1 signal to the PSTAIRE signal. Strains, +/+, JKC915; -/-, JKC1450. A and B show representatives of at least 3 biological replicates.

### Pho84 is required for ROS management

The HOG pathway is a major signaling system by which *C. albicans* induces survival responses to oxidative stress (28). We questioned whether defective HOG pathway signaling might be responsible for the hypersensitivity of *pho84* null mutant cells to neutrophil killing (29).

The phosphorylation state of the central kinase of the pathway, Hog1, was examined as a readout of HOG activation in response to oxidative stress (30). Rejecting our idea, *pho84* null mutant cells showed prolonged and hyperintense Hog1 phosphorylation during a time course of peroxide-mediated induction (Fig. 3B). This phenotype was apparent only upon exposure to extrinsic peroxide. Growing in rich medium without extrinsic oxidative stressors, *pho84* mutant, like wild type cells, exhibited a minimal signal from phosphorylated Hog1 (S3 Fig.) and we saw no difference between these strains; the Western blot may be too insensitive an assay to detect differences when signals are low.

The concurrent findings of Hog1 pathway hyperactivation of *pho84* null mutant cells, and their increased susceptibility to extrinsic ROS, suggested these cells might be unable to manage intrinsic intracellular ROS. We compared the ROS content of unstressed *pho84* null and wild type cells, and of those exposed to an oxidative stressor. In fact, cells devoid of Pho84 contained more 2’,7’-dichlorodihydrofluorescein diacetate (DCFDA)-detectable ROS, compared to the wild type, when unexposed or exposed to menadione (Figs. 4A and 4B). Their inability to manage ROS was not due to starvation for inorganic phosphate (Pi), because in Pi-replete media, *pho84* null mutant cells still contained significantly more ROS (Fig. 4A). We concluded that Pho84 is required for ROS management in *C. albicans.*

**Figure 4.**
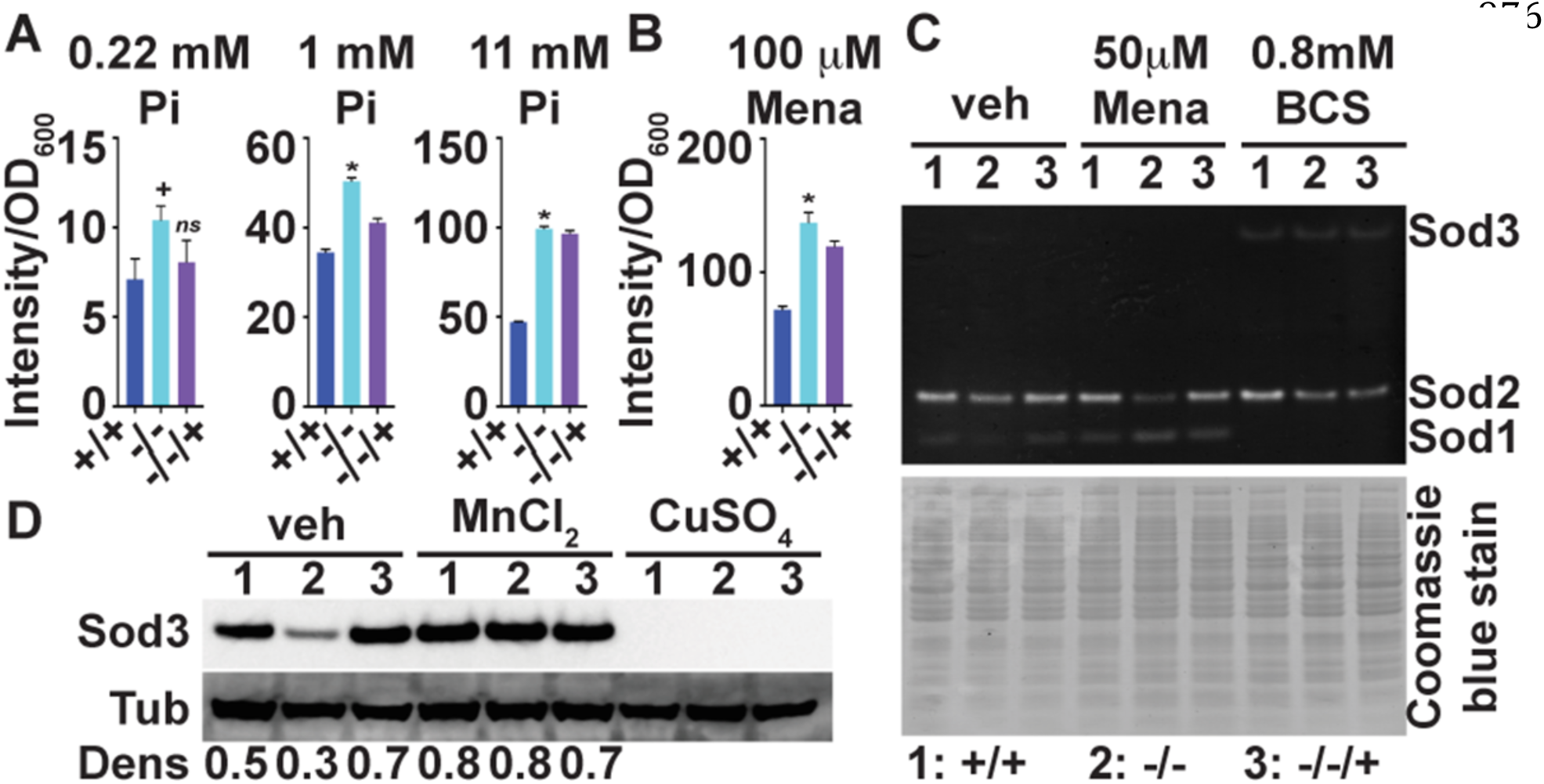
ROS management and Sod3 expression are defective in cells lacking *PHO84.* (A) DCFDA-detectable ROS of cells unexposed to extrinsic oxidative stress, of strains +/+, JKC915; -/-, JKC1450 and -/-/+, JKC1588 diluted into SC medium with 0.22 mM, 1 mM and 11 mM Pi. Fluorescence intensity was measured after staining cells with 50 μM DCFH-DA. *-/*-versus +/+ at 0.22 mM Pi *p*=0.0149. *-/-* versus +/+ at 1 mM Pi *p*<0.0001. *-/-* versus *+/+* at 11 mM Pi *p*<0.0001. *p* values per Student’s t-test. (B) DCFDA-detectable ROS production during exposure to 100 μM menadione (Mena). Strains as in A cultured overnight were diluted into SC medium (Loflo) and fluorescence intensity was measured as in A. *p*=0.0002 for *-/-* versus +/+. (C) Superoxide dismutase (SOD) activity of strains as in A, grown in YPD medium with vehicle, 50 μM Menadione (Mena) and 0.8 mM bathocuproine disulfonic acid (BCS) for 8 hours; cell lysate in non-denaturing gel stained with nitroblue tetrazolium to detect SOD activity and with Coomassie blue to assess loading. (D) Western blot of strains as in A, grown in normal SC medium with vehicle, 3 mM MnCl_2_ and 3 mM CuSO_4_ for 13 hours, probed for Sod3 and loading control tubulin. *p* values were calculated using Student’s t-test. \**p*<0.01; +*p*<0.05; ns non-significant. A-D show representatives of at least 3 biological replicates; error bars SD of 3 technical replicates.

### Superoxide dismutase expression is perturbed in *pho84* null mutant cells

Superoxide dismutases (SODs) contribute to ROS management by disproportionating the superoxide anion into H_2_O_2_ and oxygen, using a redox-active metal (15). In *C. albicans pho4* null mutants, lacking the DNA binding protein that controls the PHO regulon, mRNA expression of intracellular copper-using superoxide dismutase *SOD1* (15) is upregulated while its activity is decreased, and mRNA of manganese-using *SOD3* is decreased (31). We questioned whether increased ROS content in *pho84* null mutant cells might be due to decreased SOD protein content or -activity. *pho84* null mutant cells showed a subtle decrease of Sod1 activity during growth in standard media (Fig. 4C), as assayed by nitroblue tetrazolium (NBT) reduction in non-denaturing protein gels, which display activity of the 2 abundant intracellular SODs, Sod1 and 2 (15). During oxidative stress with menadione, variable activities of Sod1 and 2 were seen between experimental replicates, ruling out specific conclusions. Exposure to the copper chelator bathocuproine disulfonic acid (BCS) was used as a control for the identity of the bands since sparse ambient copper is known to decrease Sod1 activity (32) (Fig. 4C). Since under most conditions, Sod3 activity is not detectable on these gels, we examined its protein abundance by Western blot. Sod3 protein concentration was markedly reduced in *pho84* null mutant cells (Fig. 4D) in the absence of extrinsic oxidative stress. In the presence of high ambient manganese, Sod3 concentrations in the *pho84* mutant appeared as robust as those in the wild type (Fig. 4D). Sod3 expression dropped below the detectable limit in wild type cells in high ambient copper (Fig. 4D) as expected (32), as did that of the *pho84* null mutant. Hence, Sod3 expression is diminished in standard culture conditions in cells lacking Pho84, but can be induced in conditions of high abundance of its metal co-factor manganese.

Expression of SODs varies with metal availability in *C. albicans* (32). Since in *S. cerevisiae*, Pho84 transports manganese in addition to Pi under certain conditions (33), we questioned whether lack of *PHO84* might deplete *C. albicans* of manganese, resulting in Sod3 downmodulation. But upon measuring the intracellular manganese and copper concentrations, we found the opposite: both metals’ concentrations were increased in *pho84* null mutant cells growing in standard synthetic complete medium, compared to wild type cells (S4A and S4B Figs.). Manganese concentration in *pho84* null mutant cells supplemented with copper or manganese in the medium was like that of the wild type, while copper was increased in these cells under the same conditions (S4A and S4B Figs.). Consequently, lack of metal co-factors for SODs does not account for decreased Sod3 expression, or for slightly decreased Sod1 activity, in *pho84* null mutant cells.

### Overexpression of a TORC1 activator downstream of Pho84 suppresses ROS management defects of *pho84* null mutant cells

Since their Pi transport defect was not sufficient to explain the ROS hypersensitivity of *C. albicans* cells lacking Pho84, and since they did not exhibit lack of metal co-factors for SODs, we turned to another of their phenotypes in order to understand their ROS management defect. These cells exhibit decreased TORC1 signalling (8). This phenotype can be suppressed by overexpression of one, but not the other, of two small GTPases known to activate *C. albicans* TORC1 (8). The small GTPase Gtr1, a component of the EGO complex, which we hypothesize to participate in transmitting a Pi signal to TORC1, suppresses some *pho84* phenotypes when overexpressed (8). Not all *pho84* mutant cells’ phenotypes are suppressible in this way; e.g. *GTR1* overexpression does not suppress the hyphal morphogenesis defect of cells lacking Pho84 (not shown). We asked whether ROS management of these cells might be improved by activating TORC1 downstream of overexpressed *GTR1.* In the absence of exogenous oxidative stressors, *GTR1* overexpression decreased the DCFDA-detectable ROS in cells with active TORC1, as well as in cells experiencing TORC1 inhibition by exposure to a low concentration of rapamycin (Fig. 5A). Sod3 expression recovered in *pho84* cells overexpressing *GTR1* (Fig. 5B), but resistance to neutrophil killing did not (not shown). Previously we had shown that inhibition of Pho84 with small molecules, phosphonoacetic acid (PAA) and phosphonoformic acid (foscarnet, Fos), leads to decreased TORC1 signaling (8). To test whether small-molecule Pho84 inhibition also leads to defective ROS management, we exposed wild type cells to these compounds and measured their DCFDA-detectable ROS. Exposure to Pho84 inhibitors increased the ROS content of wild type cells unexposed to external oxidative stressors (Fig. 5C), suggesting that the ROS detoxifying role of Pho84 can be targeted pharmacologically. Together, these results indicate that activating TORC1 downstream of Pho84 can suppress a specific ROS detoxification defect of *pho84* cells, Sod3 expression, but that this is only one among the mechanisms that render these cells hypersensitive to neutrophil-imposed oxidative stress.

**Figure 5.**
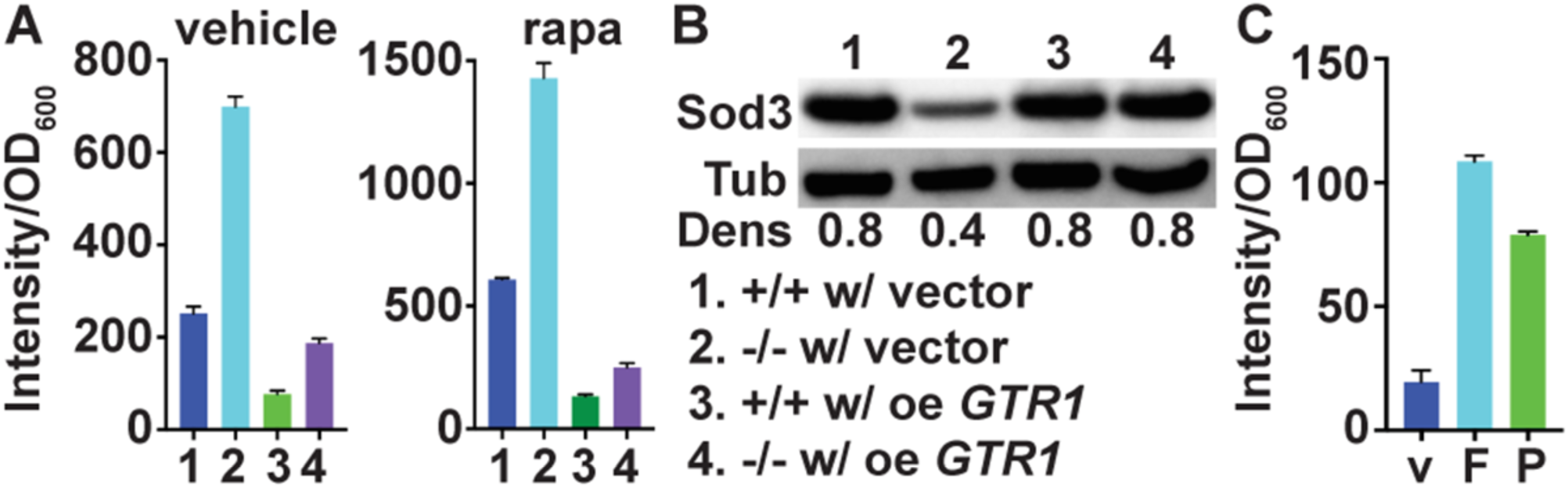
TORC1 activity contributes to ROS management and Sod3 expression. (A) DCFDA-detectable ROS measurement of strains, (1) *+/+* containing vector (JKC1594), (2) *-/*-containing vector (JKC1598), (3) *+/+* overexpressing *GTR1* (JKC1596), (4) *-/-* overexpressing *GTR1* (JKC1600). Cells cultured overnight in YPD were diluted into SC medium (Loflo) with vehicle or 8 ng/ml rapamycin for 1 hour, and fluorescence intensity was determined after staining cells with 50 μM DCFH-DA. *p*=0.0252 for ratio of *-/-* with vector and *-/-* overexpressing *GTR1* versus ratio of wild type with vector and wild type overexpressing *GTR1;* all cells exposed to vehicle. *p*=0.0014 for ratio of *-/-* with vector and *-/-* overexpressing *GTR1* versus ratio of wild type with vector and wild type overexpressing *GTR1;* all cells exposed to rapamycin. *p* values per Student’s t-test. (B) Western blot of strains as in A, grown in SC medium (Loflo) for 1 hour, probed for Sod3 and loading control tubulin. (C) Wild type (SC5314) cells were exposed to vehicle (v), 0.5 mM foscarnet (F) or 1 mM PAA (P) for 1 hour, and fluorescence intensity was determined as in A. p<0.001 for both 0.5 mM foscarnet versus vehicle and 1 mM PAA versus vehicle. A-C show representatives of at least 3 biological replicates; error bars SD of 3 technical replicates.

We then tested whether heterologously controlled overexpression of *SOD3* could directly suppress ROS hypersensitivity of cells lacking Pho84. In *PHO84* wild type and *pho84* null mutant backgrounds that express a repressible tTA (34), we replaced 50 bases of the native promoter of one *SOD3* allele with a construct in which transcription is controlled by *tetO.* In the absence of doxycyline, this construct induces high-level transcription, while it results in substantial transcriptional repression during doxycycline exposure (35). In *pho84* null mutant cells, *SOD3* overexpression largely, but not completely, suppressed in vitro sensitivity to the superoxide-inducing compound plumbagin, but it had no detectable effect in the wild type (Fig. 6). Transcriptional repression of one *SOD3* allele from *tetO* had a strong plumbagin-sensitizing effect on *pho84* null mutant cells. To control for potential artifactual effects of doxycycline exposure, we also replaced the same *SOD3* promoter sequence with a glucose-repressible *MAL2* promoter and examined the response to plumbagin. Transcriptional repression from *pMAL2* (in glucose) phenotypically resembled that from *tetO* (in doxycycline) (Fig. 6), indicating that repression of *SOD3* transcription, and not an off-target effect of doxycycline, was responsible for superoxide hypersensitivity of the cells lacking Pho84 under these conditions. Since ROS resistance was not completely recovered in *pho84* null mutant cells overexpressing *SOD3*, we concluded that misregulation of *SOD3* expression level is one, but not the only mechanism of ROS hypersensitivity of cells lacking Pho84. Cells lacking Pho84 that overexpressed *SOD3* from *tetO* did not recover resistance to whole blood or to neutrophils (not shown), indicating that in these complex ex vivo environments, Pho84 is required for further functions beyond Sod3 expression, to confer resistance to host defenses on the fungal cell.

**Figure 6.**
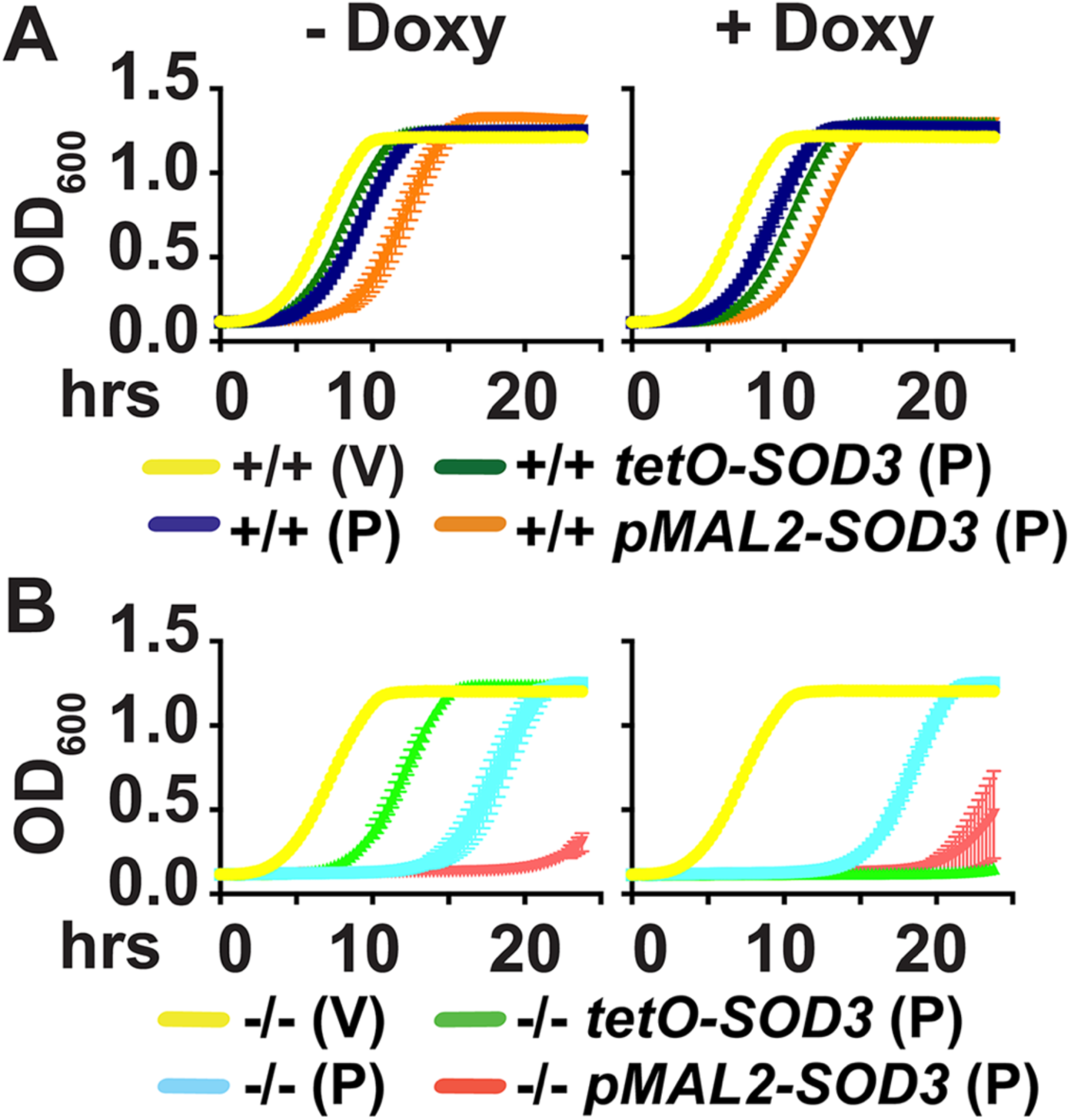
Overexpression of *SOD3* from a heterologous promoter significantly rescues ROS hypersensitivity of cells lacking Pho84. Cells were grown on YPD agar medium without or with 50 ng/ml doxycycline for 39 hrs, and were maintained in these doxycycline concentrations throughout the course of the experiment. Cells were inoculated at OD_600_ 0.1 in YPD (glucose-containing medium that represses transcription from *pMAL2)*, with vehicle DMSO (V) or 21 μM Plumbagin (P). OD_600_ was monitored every 15 minutes for strains (A) wild type background: +/+, JKC915; +/+ *tetO-SOD3/SOD3*, JKC1738; +/+ *pMAL2-SOD3/SOD3*, JKC1776; (B) *pho84* null mutant background: -/-, JKC1450; -/- *tetO-SOD3/SOD3*, JKC1745; -/- *pMAL2-SOD3/SOD3*, JKC1780. A, B are representative of 3 biological replicates, error bars SD of 3 technical replicates.

Our working model is that one mechanism, among others, by which Pho84 contributes to oxidative stress management is by activating TORC1, which in turn induces Sod3 expression (Fig. 7). This model raises several testable questions, including which downstream branch of TORC1 signaling impacts Sod3 levels, whether Sod3 is regulated at a transcriptional, translational or posttranslational level, and whether other *C. albicans* mechanisms for ROS management, affected by lack of Pho84, also depend on TORC1 signaling.

**Figure 7.**
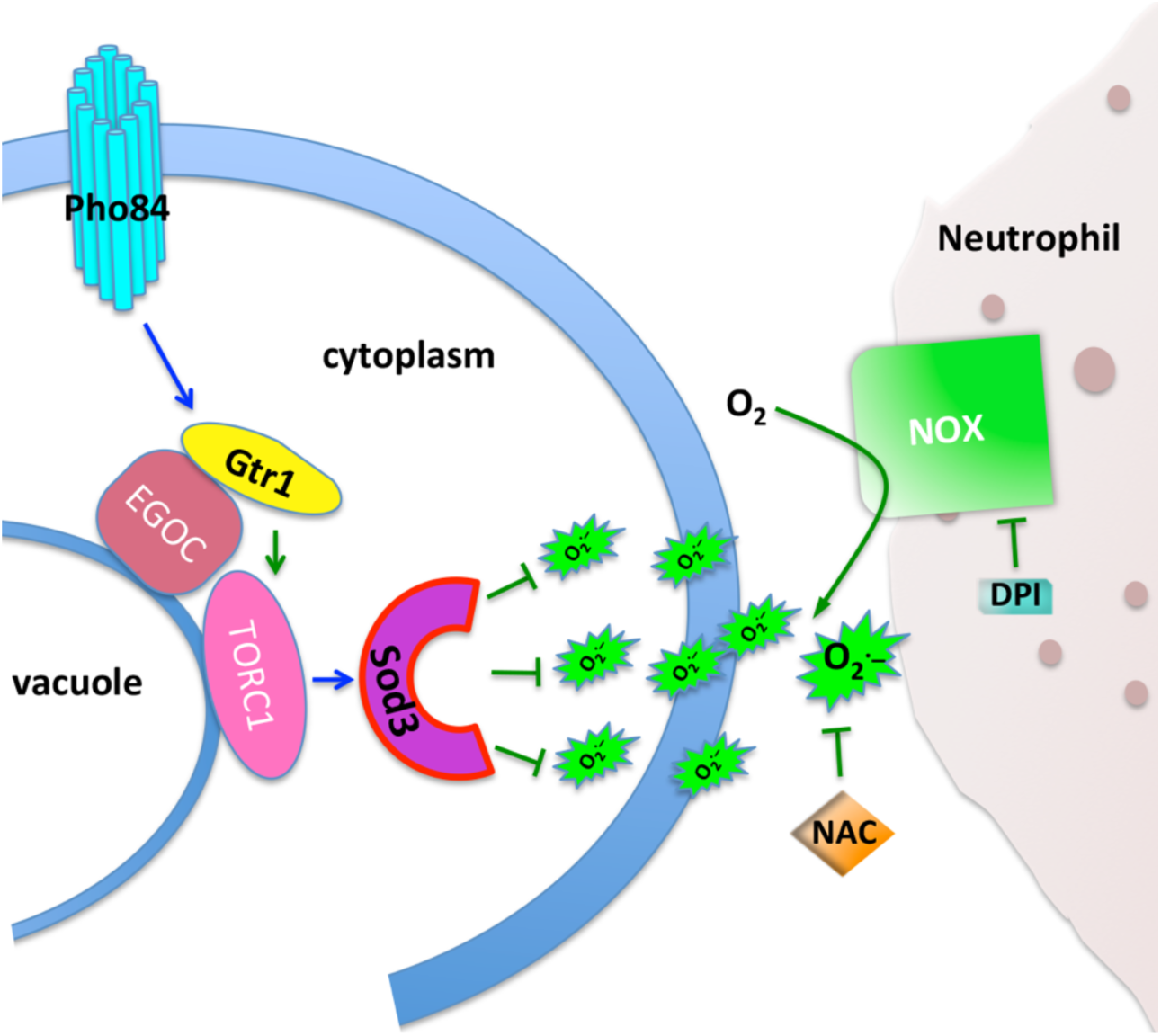
Pho84 activates TORC1 via Gtr1 to modulate Sod3 expression in combating neutrophil derived ROS. Signalling events with known molecular mechanisms in *C. albicans* are shown as green lines. Blue lines represent predicted activities based on our findings.

## Discussion

Virulence attenuation in mutants of a single macronutrient transporter like Pho84 (Fig. 1), one of 4 predicted cell-surface Pi transporters of *C. albicans*, is unusual and has not been reported to our knowledge. In addition to their hyphal growth defect (Fig. 1D), we posit that *pho84* null mutant cells’ diminished virulence is due to their hypersensitivity to neutrophil killing (Fig. 2B-D). When a neutrophil encounters an invasive pathogen, a broad array of noxious molecules and hydrolytic enzymes are exocytosed, released into the phagosome (36) or onto microbial cells during formation of extracellular traps (21). Among these, ROS are critical for killing *C. albicans* (16, 37, 38). We found that pharmacologically scavenging neutrophil-generated ROS (Fig. 2C), blocking neutrophil ROS production (Fig. 2D), or exposing *C. albicans* to neutrophils from a patient with a genetic defect in ROS production (Fig. 2E) rescued viability of *pho84* null mutant cells in this interaction. These findings indicate that among the large neutrophil armamentarium (26), *C. albicans* cells without functional Pho84 are particularly sensitive to ROS. *pho84* null mutant cells were hypersensitive in vitro to each of 3 distinct sources of exogenous ROS, plumbagin and menadione, which generate the superoxide anion, and to the peroxide source H_2_O_2_ (Fig. 3A).

In C. *albicans*, the HOG signaling pathway responds to oxidative stress to induce protective responses (28, 39). The paradox that *pho84* null mutant cells with supranormal HOG signalling exhibited increased oxidative stress sensitivity (Fig. 3) was resolved when these cells were found to harbor excessive ROS concentrations, including in unstressed normal growth conditions. This constellation of elevated intracellular ROS and hyperreactive HOG signaling suggests that *pho84* mutant cells’ capacity for detoxifying ROS is at its limit during unperturbed growth, leading to unchecked oxidative stress and increased cell death during exposure to neutrophils. Lack of sufficient Pi for NADPH synthesis, required for the ROS-detoxifying activities of glutathione- and thioredoxin reductases (40), could also account for ROS hypersensitivity of *pho84* mutant cells. However, in high ambient Pi, where the contribution of Pho84 to intracellular Pi stores is less critical (8), their ROS were still ~2-fold those of the wild type (Fig. 4A), evincing an additional mechanism.

The first line of defense against exogenous ROS released by host phagocytes onto a fungal cell are SODs bound to the cell surface (41, 42). Their importance for *C. albicans* is underscored by an unusually numerous complement of 3 extracellular SODs in this species (15). Among extracellular SODs, hyphal-specific Sod5 was recently shown to participate in morphogenetic regulation by disproportionating superoxide anion generated by the newly discovered *C. albicans* NADPH oxidase Fre8, since the resulting peroxide stimulates hyphal extension (43). Transcription of *SOD5* is upregulated in cells lacking the PHO regulon transcription factor Pho4, which induces *PHO84* expression (31), and whose nuclear localization is predicted to be induced in the absence of Pho84 (44). Potential relationships between Pho4-regulated *PHO84* and *SOD5* expression and activities, and their possible effect on hyphal morphogenesis, await further experimental examination.

If exogenous superoxide anion escapes detoxification by extracellular SODs and becomes protonated, it can cross the cytoplasmic membrane to damage intracellular macromolecules; cytosolic SODs are present to scavenge these ROS (15). We found a sharp decrease in the protein level of the cytosolic superoxide dismutase Sod3 in *pho84* null mutant cells. Sod3 is a singular manganese-requiring cytosolic SOD (45), one of an unusual pair of cytosolic SODs encoded only by some fungal genomes (32). As during invasive infection *C. albicans* cells are exposed to drastic changes in copper concentrations, they switch between expression of cytosolic copper-Sod1 during copper repletion or manganese-Sod3 during copper starvation (32). Expression of *PHO84* is controlled by the Pho4 transcription factor, which regulates almost a thousand genes in response to Pi availability (31). The contribution of *C. albicans* Pho4 to virulence is attributed to its role in metal homeostasis (31). In contrast, we found Pho84 to be dispensable for metal repletion of the cell (Fig. S3).

Failing to find a defect in metal uptake of *pho84* null mutant cells to account for decreased Sod3 levels (Fig. S3), we looked to their defect in TORC1 signalling. The mitochondria are major sources of ROS which are byproducts of respiration. TORC1 and mitochondrial ROS both impact the yeast chronological lifespan and aging (46), and mitochondrial protein quality control affects TORC1 signaling and *C. albicans* virulence (47). But a requirement for active TORC1 to induce cytosolic SOD expression and thereby manage innate or exogenous ROS (Fig. 5) has not previously been described in any organism to our knowledge. Further work will have to determine whether the TORC1-activating function of Pho84 depends on Pi transport or whether it resides in a genetically or chemically separable signalling activity, like the transceptor activity of *S. cerevisiae* Pho84 towards PKA (48, 49). Our finding that Sod3 protein levels are decreased in cells lacking Pho84, but can be recovered when TORC1 is activated downstream of Pho84 (Fig. 5), opens further areas of investigation. Among them is the relationship between TORC1 activity and that of other SODs, since one could speculate that active TORC1 corresponds to an active metabolic state, which gives rise to increased intrinsic ROS and requires increased SOD activity.

TORC1 signaling integrates information and responses to a large number of parameters, most of which are likely to be involved in the interaction with competing members of the microbiome and with the host. Hence we do not expect that simply upregulating TORC1 by overexpressing *GTR1* enhances *C. albicans* virulence. It is likely the flexibility, not the intensity of pro-anabolic TORC1 signaling that maximizes fungal survival and proliferation. Our findings nevertheless highlight the importance of Pi signaling as one input to TORC1 for the capacity of the fungus to withstand stressors imposed by the host and maximize its virulence potential.

Resistance of *C. albicans pho84* null mutant cells to in vitro ROS exposure was significantly restored by overexpression of *SOD3* (Fig. 6). Transcriptional repression of one *SOD3* allele resulted in synthetic ROS hypersensitivity with deletion of *PHO84* (Fig. 6). However, resistance to ex vivo stressors, whole blood and neutrophil killing, was not restored to *pho84* mutant cells by overexpression of *SOD3* from the heterologous *tetO* construct. Hence, maintenance of Sod3 levels via TORC1 is only part of the mechanisms by which Pho84 contributes to resistance to host defenses. Other mechanisms, including provision of Pi for NADPH-dependent ROS management and interactions with other SODs, await experimental identification.

In addition to reactive oxygen species detoxification, Pho84 also contributes to hyphal growth of *C. albicans* (8). In the murine disseminated candidiasis model, we observed reduced hyphal growth of cells lacking Pho84 in vivo. A requirement for SOD activity in virulence was found each time it was examined (20, 41, 50); this is not the case for hyphal growth (51-53). Nevertheless, defective regulation of hyphal morphogenesis may also contribute to *pho84* mutants’ virulence attenuation. This defect is not suppressed by *GTR1* overexpression. Loss of Pho84 appears to affect virulence of *C. albicans* by further mechanisms beyond those we have defined so far.

We previously showed that *C. albicans* Pho84 and its TORC1-activating function can be inhibited by small molecules (8), raising the possibility of identifying further inhibitors with favorable therapeutic indices. The compounds we investigated include an FDA-approved antiviral agent, which at concentrations achieved in plasma during therapeutic dosing inhibits hyphal morphogenesis and potentiates the in vitro activity of important medications representing 2 of the 3 classes of antifungal agents in clinical use (8). Pho84 has no human homolog, so its inhibitors can spare human targets. We now find that these compounds also perturb ROS management (Fig. 5C), so that inhibiting Pho84 with novel compounds could simultaneously potentiate antifungal drugs and impair *C. albicans* virulence.

Pathogenic unicellular parasites require intact Pi homeostasis for development and virulence (9, 11, 14, 54), as do bacterial pathogens (55). Among fungi, representatives of all extant lineages possess core components of the PHO regulon, including high-affinity H+-Pi symporters like Pho84 (56). In a basal fungal phylum, the water mold *Blastocladiella emersonii’s* complex life cycle (57) depends on Pi availability (56, 58) as it progresses between flagellated zoospores, their germination, vegetative growth with production of a rhizoidal system, and sporulation (58). Pi uptake and expression of PHO regulon components correspond to the life cycle stage of this fungus, indicating an ancient origin of the PHO system at or near the emergence of the fungal kingdom and suggesting its early connection to developmental regulation (56). Links of Pi homeostasis with the complex developmental steps required in *C. albicans* pathogenesis hence have primordial precedents in its primitive fungal predecessors. Advancing analysis of Pi acquisition and homeostasis in fungal pathogens (31, 59-63) to the level of understanding achieved in model fungi like *S. cerevisiae, Neurospora crassa* and mycorrhizal fungi (44, 64-68), could elucidate further virulence determinants and, given fundamental differences between human and fungal phosphate homeostasis, potentially identify much-needed further targets of antifungal therapy.

## Methods

### Ethics Statement

For experiments utilizing human neutrophils whose donors were identifiable, written informed consent was obtained from study subjects, according to protocols approved by the Boston Children’s Hospital Institutional Review Board. For human neutrophils isolated from anonymized blood bank donation aliquots, no informed consent was obtained because the donors were not identifiable, and this procedure was approved by the Boston Children’s Hospital Institutional Review Board. The Blood Bank Laboratory at the Boston Children’s Hospital provided the anonymized blood samples.

All animal experiments were performed in accordance with recommendations in the Guide for the Care and Use of Laboratory Animals of the National Institutes of Health. The Institutional Animal Care and Use Committee of Los Angeles Biomedical Research Institute at Harbor-UCLA Medical Center, Torrance, CA reviewed and approved the murine oropharyngeal candidiasis experimental protocol, IACUC Project No. 30842-01 (A3330-01). All animal experimentation of the intravenously infected, disseminated murine candidiasis model was conducted in an AAALAC-certified facility at The University of Texas at San Antonio (UTSA) following the National Institutes of Health guidelines for housing and care of laboratory animals and performed in accordance with institutional regulations after pertinent review and approval by the Institutional Animal Care and Use Committee at The University of Texas at San Antonio. UTSA Animal Welfare Assurance ID D16-00357 and UTSA IACUC Protocol Number MU018. Mice were allowed a 1-week acclimatisation period before experiments were started.

### Strains and culture conditions

*C. albicans* strains used are shown in Table 1. Strains were constructed as in (8); introduced mutations were confirmed by PCR spanning the upstream and downstream homologous recombination junctions of transforming constructs, and sequencing. Only prototrophs were compared in each experiment. At least two strains of newly constructed genotypes were examined phenotypically. Only after confirming that their phenotypes were the same, was one strain of each genotype chosen for representation. Experiments with defined ambient Pi concentrations were performed in YNB 0 Pi (ForMedium Ltd, Norfolk, UK) with added KH2PO4 to stated concentrations. Low fluorescence medium for ROS measurement was made from YNB Loflo (ForMedium Ltd, Norfolk, UK). Other media were used as previously described (35).

Culture conditions of cells during growth before each experiment, e.g. overnight, were standardized for each set of experiments. Normal saline (0.9% NaCl) was used for all wash steps to avoid uncontrolled exposure to Pi in PBS. Cells used for *SOD3* repression or - overexpression experiments were maintained in conditions without or with 50 ng/ml doxycycline or using maltose or glucose as carbon source, throughout the course of the experiment, including during washes, beginning at the point of reviving frozen stocks.

Nomenclature of strains used in text and figure legends is +/+ *(PHO84/PHO84*, wild type); -/- *(pho84/pho84*, null mutant) and -/-/+ *(pho84/pho84::PHO84*, reintegrant of intact *PHO84* at the native locus).

### Growth Assays

For growth curves in liquid media, cells grown on a YPD plate for 2 days were washed once in 0.9% NaCl and diluted to an OD_600_ of 0.1 in 150 μl medium in flat bottom 96-well dishes. OD_600_ readings were obtained every 15 min in a BioTek™ Synergy™ 2 Multi-Mode Microplate Reader (Winooski, VT, USA).

At least 3 biological replicates were obtained on different days. Standard deviations of 3 technical replicates, representing separate wells, were calculated and graphed in Graphpad Prism.

### Western Blots

Cell harvesting, lysis and Western blotting were performed as described in (69). Antibodies used are listed in Table 4. At least three biological replicates were obtained on different days. For densitometry, ImageJ (imagej.net/welcome) software (opensource) was used to quantitate signals obtained on a KODAK Image Station 4000MM.

### Infection of flies

The wild-type *Drosophila melanogaster* OrR strain obtained from the Bloomington stock center was maintained on standard cornmeal agar medium at 25°C. Single colonies of *C. albicans* strains on YPD were used to inoculate overnight cultures at 30°C. The cultures were diluted in fresh YPD to an OD_600_ of 0.1 and incubated shaking at 30°C until an OD_600_ of 1.0. A 1 ml aliquot of the culture was centrifuged at 16,000 x g, washed once in phosphate-buffered saline (PBS; pH 7) and re-suspended in 1 ml of PBS.

For infection of flies, protocols were followed according to (17). Male and female flies, 1-5 days old, were injected with approximately 50 nl fungal cell suspensions (approx. 500 cells/fly) using a fine glass capillary needle with a micro-injector (TriTech Research, Los Angeles, CA, USA). Cohorts of 30 flies were injected and maintained in separate vials. At least 3 (up to 7) biological replicates (independently prepared fungal preparations derived from individual colonies) and 6 -15 technical replicates per strain were carried out. Infected flies were maintained at 29°C for up to five days after infection and the number of surviving flies was noted on daily basis.

### Oropharyngeal candidiasis

Mice were infected by inoculating with *Candida albicans* cells sublingually as in (18). Eight mice were infected for each *C. albicans* strain. Fungal burden was calculated by counting CFU per gram tissue on agar medium after homogenizing tongue tissue.

### Murine model of hematogeneously disseminated candidiasis

Cultures of *C. albicans* strains for injection were grown overnight in YPD medium and incubated at 30°C. *C. albicans* cells (4 × 10^5^) were delivered by tail vein injection into mice, each consisting of eight 6-to-8-week-old female BALB/c mice. Days on which mice died were recorded, and moribund animals were euthanised and recorded as dying the following day.

For histology, kidneys retrieved from sacrificed mice were fixed in 10% buffered formalin and embedded in paraffin, and thin tissue slices were obtained and stained with Grocott-Gomori Methenamine Silver (GMS) stain prior to microscopic evaluation.

### Whole blood killing assay

Peripheral blood was collected by venipuncture of healthy adults who had not taken any medications during the ten days prior to the experiment. Blood was gently drawn through a 21-gauge needle into heparinised glass vacutainers. Before exposure to blood or other stressors, the *C. albicans* cells were inoculated into 10 ml YPD at a density of OD_600_=0.2, and incubated for 15 hours at 30°C with shaking at 200 rpm. Harvested cells were washed once with normal saline and diluted to a concentration of 5×10^4^/ml counted by an Innovatis CASY Cell Counter (Roche). Fifteen μl cells were added into a 96-well plate first (VWR, Cat. # 62406-081) and incubated with 135 μl blood for indicated time points. To determine CFU/ml, cells were spread on Trypticase Soy Agar with 5% Sheep Blood (Thermofisher Scientific, Cat. # B21239X) or on laboratory-made YPD plates and colonies were counted after incubation at room temperature for 2 days. At least 3 biological replicates were performed on different days for each result shown.

### Isolation of human neutrophils

Human primary neutrophils were isolated from apheresis-derived buffy coats provided by the Blood Bank Laboratory at the Boston Children’s Hospital. Erythrocytes were sedimented by adding an equal volume of dextran/saline solution (3% dextran in PBS) at room temperature for 30 min. The erythrocyte-depleted supernatants were then layered on Lymphocyte Separation Medium (Lymphoprep, axis-shield) and centrifuged at 400 g at room temperature for 30 min. Contaminating erythrocytes in the neutrophil pellets were lysed by a brief (<30 second) treatment with ice cold water and 0.6 M KCl. The purity of neutrophils was >90% as determined by Wright–Giemsa staining.

Peripheral blood was collected from healthy adult volunteers or from a CGD patient into heparinized glass vacutainers. Granulocytes were isolated using Polymorphoprep (axis-shield) according to manufacturer instructions. Briefly, whole blood was loaded on top of a Polymorphoprep gradient and centrifuged at 500g for 30 min without brake. According to manufacturer instructions, the neutrophil containing band was harvested. Osmolarity was reestablished by adding an equal volume of 50% HBSS and neutrophils were pelleted. Contaminating red blood cells were lysed by treating the pellet twice with 9 ml of cell culture ready dH_2_O (Gibco A1287301) and reestablishing osmolarity with 1 ml of 10x PBS. Neutrophils were obtained at a purity >95% with the contaminating cells mainly being RBCs. Boston Children’s Hospital Institutional Review Board protocols were followed for all human blood samples.

### Neutrophil killing assay

*C. albicans* strains were incubated with HL-60 derived neutrophils for 90 minutes at 37°C with 5% CO_2_ and quantitative culturing on Trypticase Soy Agar with 5% Sheep Blood plates (Thermofisher Scientific, Cat. # B21239X). Percent survival was calculated by dividing the number of CFU after co-culturing with HL-60 derived neutrophils by the number of CFU from *C. albicans* incubated with media without neutrophils. HL-60 derived neutrophils were tested at a 20:1 phagocyte:fungus ratio in RPMI plus 10% pooled human serum.

*C. albicans* strains were incubated with freshly isolated human neutrophils for the indicated time points at 37°C with 5% CO_2_. To determine CFU/ml, cells were spread on Trypticase Soy Agar with 5% Sheep Blood (Thermofisher Scientific, Cat. # B21239X) or on laboratory-made YPD plates and colonies were counted after incubation at room temperature for 2 days.

Isolated human neutrophils were tested in HBSS medium (GIBCO, Cat. # 14025076). To scavenge ROS or inhibit ROS formation, neutrophils were treated with N-acetyl-l-cysteine (NAC) (Sigma) for 90 min at M.O.I. 2, or pretreated with 10μM Diphenyleneiodonium, DPI, (Tocris) for 5 minutes.

At least 3 biological replicates were obtained for each experiment on different days.

### Phagocytosis assay

*C. albicans* cells were grown overnight for 15 hours as described above and washed once with HBSS. 1 × 10^5^ Neutrophils were incubated with *C. albicans* yeast cells expressing GFP at M.O.I. 2 and 10 for 30 min. Phagocytosing neutrophils were quantified as CD11b+ GFP+ Cells. Two biological replicates were obtained on different days.

### Neutrophil ROS production assay

Extracellular ROS production was determined as described in (70). Briefly, neutrophils were stimulated with *C. albicans* yeast at M.O.I. 2 for 1 hour in the presence of 100mM Cytochrome C. To measure superoxide release, the difference in absorbance of the oxidated form of Cytochrome C (550 nm) and the background absorbance (540 nm) was measured every two minutes in a FLUOstar Omega Microplate Reader. To measure intracellular ROS, neutrophils were stimulated with *C. albicans* yeast at M.O.I. 2 for 30 minutes and subsequently stained with the ROS sensitive dye chloromethyl-H2DCFDA (Thermofisher Scientific, Cat. # D399) for 30 min. Two biological replicates were obtained on different days.

### ROS measurement in *Candida albicans*

Yeast cells grown overnight in YPD for 15 hours were washed twice with normal saline and diluted into SC medium (Loflo) supplemented with different concentrations of KH2PO4 at OD_600_ 0.5. The fluorescent dye 2’,7’-dichlorodihydrofluorescein diacetate (DCFDA) (Sigma, Cat# D6883) was added into the medium to a final concentration of 50 μM. After incubation for 50 min, cells were washed twice with normal saline. The intensity of fluorescence was read in a TE-CAN plate reader at excitation-485 nm and emission wavelength 528 nm (71), and a ratio of intensity with OD_600_ of the culture was calculated. At least 3 biological replicates were obtained for each experiment on different days.

### SOD gel activity assay and immunoblot

Cells were grown for 13 hours in SC medium with starting OD_600_ 0.01. Harvested cells were washed with sterile MilliQ H_2_O once and lysed by bead beating. A 10% Tris-glycine gel (Invitrogen, Cat. # XP00100BOX) gel was used for running samples and stained with nitroblue tetrazolium as described in (72). For Western blotting, the membrane was probed with primary antibodies: anti-Sod3 antibody (72) and anti-tubulin rat monoclonal Ab (Abcam, Cat. # ab6161) as loading control. Secondary antibodies were anti-rabbit secondary antibody (Santa Cruz Biotechnology, Cat. # 2370) and anti-rat secondary antibody (Santa Cruz Biotechnology, Cat. #97057). The blots were imaged using a KODAK Image Station 4000MM. At least 3 biological replicates were obtained for each experiment on different days.

### Intracellular metal measurement

Cells were grown in SC medium with starting OD_600_=0.01 for 13 hours. Harvested cell were washed twice with cold 1X TE (10 mM Tris, 1 mM EDTA pH 7.5) and once with cold milliQ water. Cells were resuspended in 1 mL of deionised water and 10 OD_600_ cells were used for analysis of Cu and Mn by a PerkinElmer Life Sciences AAnalyst 600 graphite furnace atomic absorption spectrometer. Three biological replicates were obtained for each experiment on different days.

### Statistical analysis

Statistical analysis was performed by Kaplan-Meier analysis for survival curves, and unpaired Student’s t test for comparison of numerical values in Prism 7 Graphpad software (GraphPad Software, Inc., CA, USA).

## Acknowledgments

We gratefully acknowledge expert performance of murine disseminated candidiasis experiments by Anna L. Lazzell. We thank Matthew Pettengill, David Dowling and Ashraf Ibrahim for helpful conversations and protocols. We thank Simon Dove for discussions and critical reading of the manuscript.

## Supporting Information

**S1 Figure.**
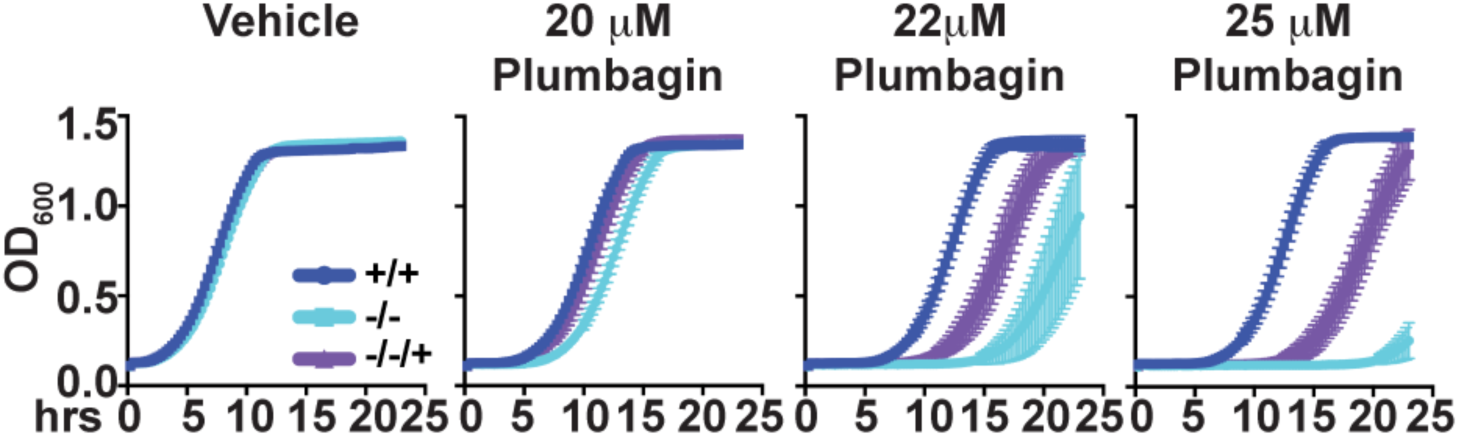
Stress intensity-dependent haploinsufficiency of *pho84/pho84::PHO84* reintegrant cells. Overnight cultures were washed with normal saline and diluted to YPD with vehicle, 20 μM plumbagin, 22 μM plumbagin and 25 μM plumbagin to OD_600_=0.1. OD_600_ was monitored every 15 minutes. Strains used were: +/+, JKC915; -/-, JKC1450 and -/-/+, JKC1588. 25 μM plumbagin depicts the same experiment as that shown in Fig. 3A. Representative of 3 biological replicates, error bars show SD of 3 technical replicates.

**S2 Figure.**
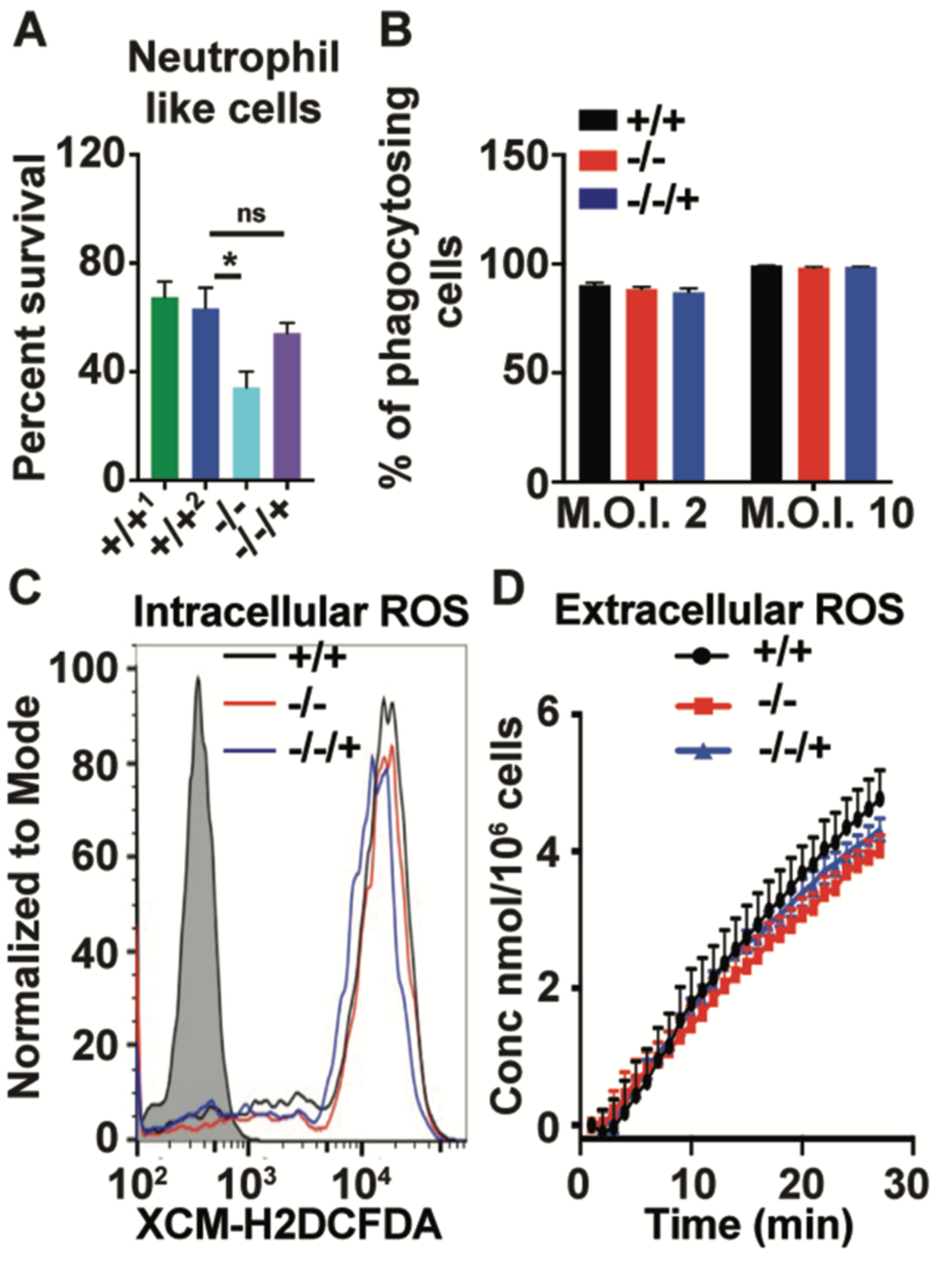
Pho84 does not affect induction of phagocytosis or ROS production by neutrophils. (A) Percent survival of *C. albicans* cells after incubation of HL-60 derived neutrophils with +/+^1^, SC5314; +/+^2^, JKC915; -/-, JKC1450 and -/-/+, JKC1500 at a 20:1 phagocyte: fungus ratio. \**p*<0.01; ns non-significant. (B) *+/+ (PHO84/PHO84 _promoter_ACT1-GFP)*, JKC1648; -/- *(pho84/pho84 _promoter_ACT1-GFP)*, JKC1651 and -/-/+ *(pho84/pho84::PHO84 _promoter_ACT1-GFP)*, JKC1653 cells were incubated with neutrophils at M.O.I. 2 and M.O.I. 10. Phagocytosing neutrophils were quantified as CD11b+ GFP+ Cells. (C) Intracellular ROS production by neutrophils was measured after stimulation with *C. albicans* yeast at M.O.I. 2 for 30 minutes. (D) Extracellular ROS production was measured by incubation with *C. albicans* yeast at M.O.I. 2 for 1 hour in the presence of 100mM Cytochrome C. Strains used were: +/+, JKC915; -/-, JKC1450 and -/-/+, JKC1588. A shows representative of 3 biological replicates; B, C: representative of 2 biological replicates is shown; error bars SD of 3 technical replicates.

**S3 Figure.**
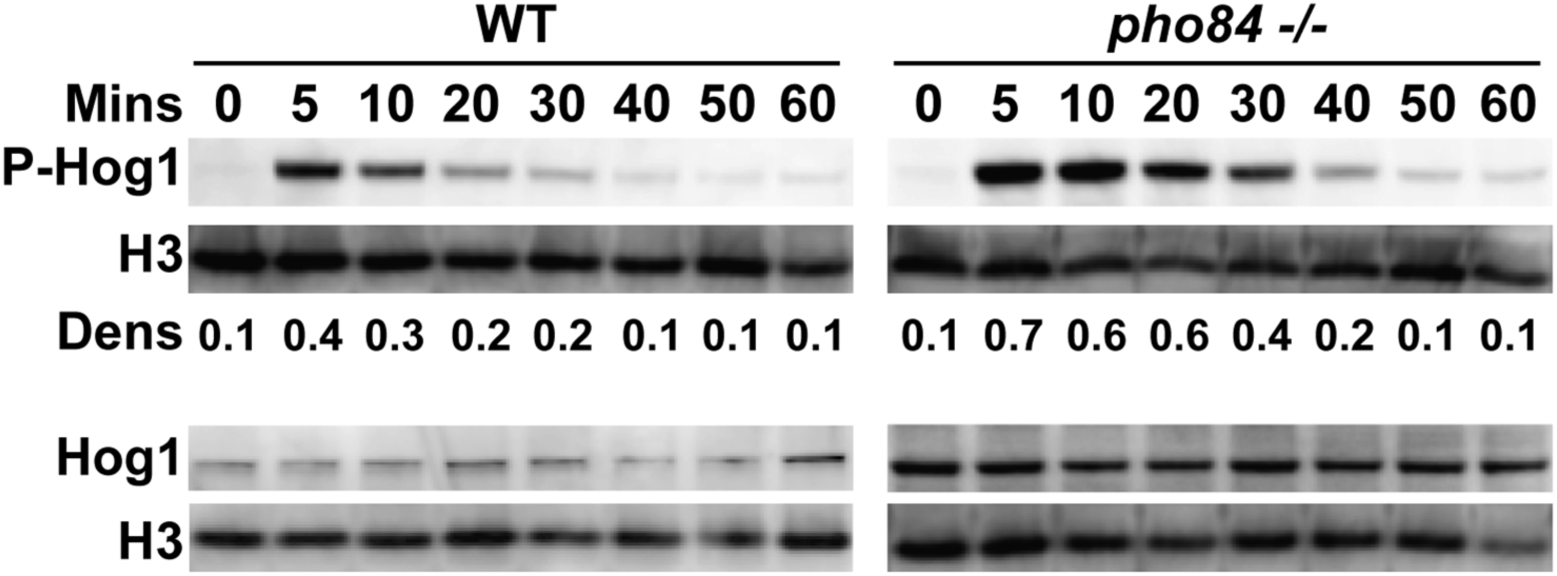
In unstressed *pho84* cells, Hog1 phosphorylation is not measurably increased, while upon extrinsic peroxide exposure P-Hog1 increase is more pronounced and prolonged in *pho84* cells than in wild type. Western blot of cells incubated in YPD without H_2_O_2_ (0 minutes) and in YPD containing 5mM H_2_O_2_ for the stated times. The membrane was probed for phosphorylated Hog1 (P-Hog1), total Hog1, and loading control Histone H3. Dens: ratio of densitometry of P-Hog1 signal to H3 signal. Strains, +/+, JKC915; -/-, JKC1450.

**S4 Figure.**
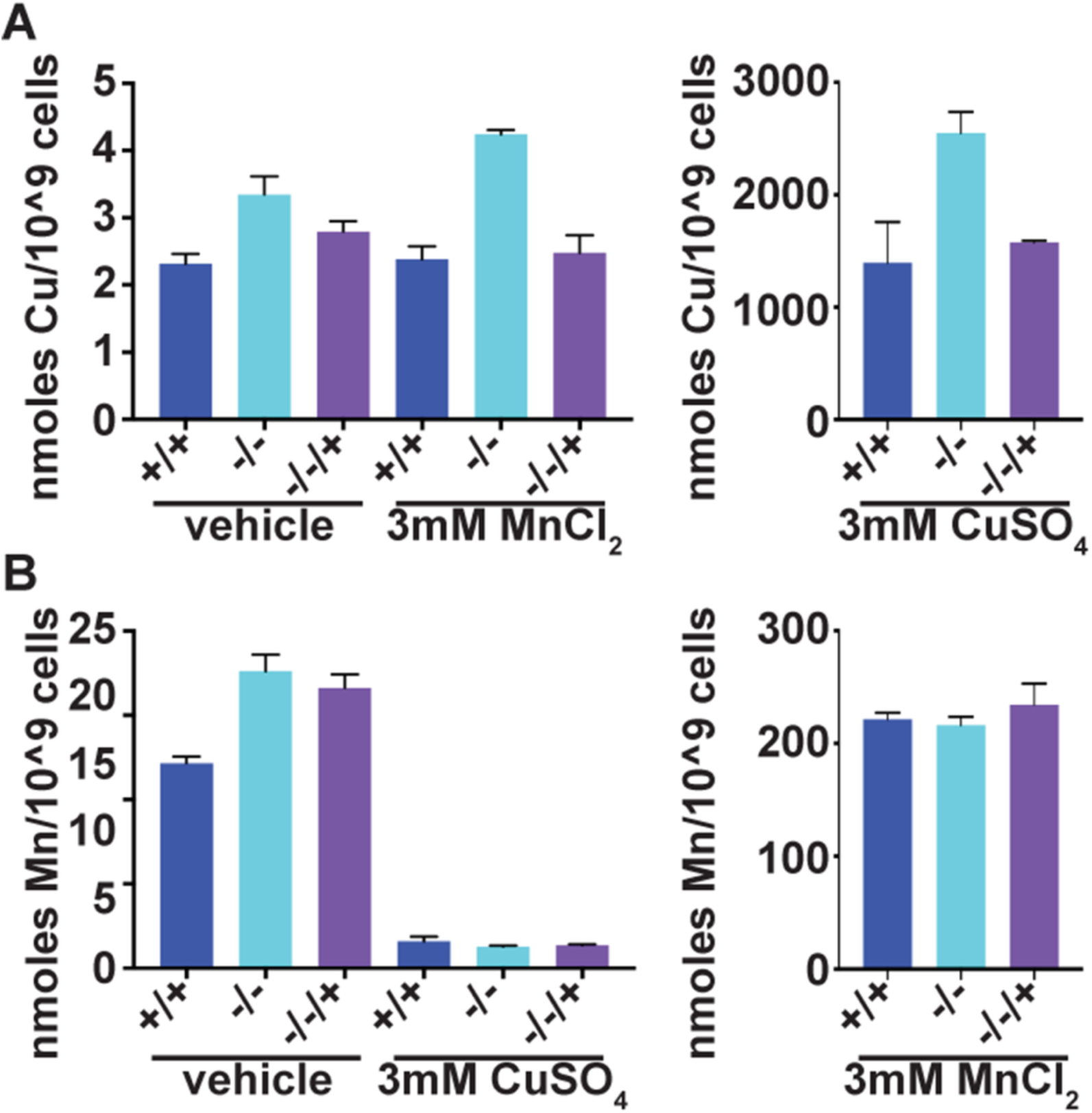
Total intracellular metal SOD co-factors are not diminished in cells lacking *PHO84.* (A) Total intracellular copper of strains +/+, JKC915; -/-, JKC1450 and -/-/+, JKC1588, grown in normal SC medium with vehicle, 3 mM MnCl_2_ and 3 mM CuSO_4_ for 13 hours was measured by atomic absorption spectroscopy (AAS). (B) Total intracellular manganese of strains as in A, grown as in A, was measured by AAS. A and B show mean of 3 biological replicates obtained on different days; error bars SD of 3 biological replicates.

**S1 Table.** Strains used in this study

| **C. albicans strain name** | **Parent** | **Genotype** | **Strain background /construction** | **Reference** |
| --- | --- | --- | --- | --- |
| SC5314 |  | Wild type | Bloodstream isolate | (1) |
| JKC915 | SC5314 | *HIS1/his1::tetR-FRT* |  | (2) |
| JKC1450 | JKC1423 | *pho84::HIS1/pho84::ARG4*<br>*his1/his1::tetR-FRT arg4/arg4*<br>*IRO1/iro1Δ::λimm<sup>434</sup> URA3/ura3Δ::λimm<sup>434</sup>* |  | (3) |
| JKC1583 | JKC1423 | *PHO84/pho84::HIS1*<br>*his1/his1::tetR-FRT arg4/arg4::ARG4*<br>*IRO1/iro1Δ::λimm<sup>434</sup> URA3/ura3Δ::λimm<sup>434</sup>* |  | (3) |
| JKC1500 | JKC1450 | *PHO84-FRT/pho84::ARG4*<br>*his1/his1::tetR-FRT arg4/arg4*<br>*URA3/ura3::λimm<sup>434</sup> IRO1/iro1::λimm<sup>434</sup>* |  | (3) |
| JKC1588 | JKC1500 | *PHO84-FRT/pho84::ARG4 LEU2/leu2::C.d. HIS1*<br>*his1/his1::tetR-FRT arg4/arg4*<br>*URA3/ura3::λimm<sup>434</sup> IRO1/iro1Δ::λimm<sup>434</sup>* |  | (3) |
| JKC1594 | JKC917 | *ACT1/<sub>promoter</sub>ACT1-NAT1-terminatorACT1*<br>*his1/his1::tetR-FRT arg4/arg4*<br>*IRO1/iro1Δ::λimm<sup>434</sup> URA3/ura3Δ::λimm<sup>434</sup>* |  | (3) |
| JKC1596 | JKC917 | *ACT1/<sub>promoter</sub>ACT1-GTR1-NAT1-terminatorACT1*<br>*his1/his1::tetR-FRT arg4/arg4*<br>*IRO1/iro1Δ::λimm<sup>434</sup> URA3/ura3Δ::λimm<sup>434</sup>* |  | (3) |
| JKC1598 | JKC1450 | *ACT1/<sub>promoter</sub>ACT1-NAT1-terminatorACT1*<br>*pho84::HIS1/pho84::ARG4*<br>*his1/his1::tetR-FRT arg4/arg4 IRO1/iro1Δ::λimm<sup>434</sup>*<br>*URA3/ura3Δ::λimm<sup>434</sup>* |  | (3) |
| JKC1600 | JKC1450 | *ACT1/<sub>promoter</sub>ACT1-GTR1-NAT1-terminatorACT1*<br>*pho84::HIS1/pho84::ARG4*<br>*his1/his1::tetR-FRT arg4/arg4 IRO1/iro1Δ::λimm<sup>434</sup>*<br>*URA3/ura3Δ::λimm<sup>434</sup>* |  | (3) |
| JKC1648 | JKC915 | *ACT1/<sub>promoter</sub>ACT1-GFP-NAT1-terminatorACT1*<br>*his1/his1::tetR-FRT* | JKC915 transformed with pJK1323, cut with BsrGI, to overexpress GFP | This work |
| JKC1651 | JKC1450 | *ACT1/<sub>promoter</sub>ACT1-GFP-NAT1-terminatorACT1*<br>*pho84::HIS1/pho84::ARG4*<br>*his1/his1::tetR-FRT arg4/arg4 IRO1/iro1Δ::λimm<sup>434</sup>*<br>*URA3/ura3Δ::λimm<sup>434</sup>* | JKC1450 transformed with pJK1323, cut with BsrGI, to overexpress GFP | This work |
| JKC1653 | JKC1588 | *ACT1/<sub>promoter</sub>ACT1-GFP-NAT1-terminatorACT1*<br>*PHO84-FRT/pho84::ARG4 LEU2/leu2::C.d. HIS1*<br>*his1/his1::tetR-FRT arg4/arg4*<br>*URA3/ura3::λimm<sup>434</sup> IRO1/iro1Δ::λimm<sup>434</sup>* | JKC1588 transformed with pJK1323, cut with BsrGI, to overexpress GFP | This work |
| JKC1361 | JKC917 | *his1/his1::tetR-FRT arg4/ARG4*<br>*IRO1/iro1Δ::λimm<sup>434</sup> URA3/ura3Δ::λimm<sup>434</sup>* |  | (3) |
| JKC1717 | JKC915 | *HIS1/his1::tetR-FRT SOD3/FLP-NAT-tetO-SOD3* | JKC915 transformed with pJK1351, cut with KpnI/NcoI, to control *SOD3* expression from *tetO*, isolate #3 | This work |
| JKC1738 | JKC1717 | *HIS1/his1::tetR-FRT SOD3/FRT-tetO-SOD3* | JKC1717 after recombination of FRTs following FLP induction | This work |
| JKC1720 | JKC915 | *HIS1/his1::tetR-FRT SOD3/FLP-NAT-tetO-SOD3* | JKC915 transformed with pJK1351, cut with KpnI/NcoI, to control *SOD3* expression from *tetO*, isolate #4 | This work |
| JKC1742 | JKC1720 | *HIS1/his1::tetR-FRT SOD3/FRT-tetO-SOD3* | JKC1720 after recombination of FRTs following FLP induction | This work |
| JKC1726 | JKC1450 | *pho84::ARG4/pho84::HIS1 SOD3/FLP-NAT-tetO-SOD3* | JKC1450 transformed with pJK1351, cut with KpnI/NcoI, to control *SOD3* expression from *tetO*, isolate #5 | This work |
| JKC1745 | JKC1726 | *pho84::ARG4/pho84::HIS1 SOD3/FRT-tetO-SOD3* | JKC1726 after recombination of FRTs following FLP induction | This work |
| JKC1729 | JKC1450 | *pho84::ARG4/pho84::HIS1 SOD3/FLP-NAT-tetO-SOD3* | JKC1450 transformed with pJK1351, cut with KpnI/NcoI, to control *SOD3* expression from *tetO*, isolate #7 | This work |
| JKC1751 | JKC1729 | *pho84::ARG4/pho84::HIS1 SOD3/FRT-tetO-SOD3* | JKC1729 after recombination of FRTs following FLP induction | This work |
| JKC1769 | JKC915 | *HIS1/his1::tetR-FRT SOD3/FLP-NAT-<sub>promoter</sub>MAL2-SOD3* | JKC915 transformed with pJK1353, cut with KpnI/NcoI, to control *SOD3* expression from *promoter*MAL2, isolate #3 | This work |
| JKC1776 | JKC1769 | *HIS1/his1::tetR-FRT SOD3/FRT-<sub>promoter</sub>MAL2-SOD3* | JKC1769 after recombination of FRTs following FLP induction | This work |
| JKC1772 | JKC1450 | *pho84::ARG4/pho84::HIS1 SOD3/FLP-NAT-<sub>promoter</sub>MAL2-SOD3* | JKC1450 transformed with pJK1353, cut with KpnI/NcoI, to control *SOD3* expression from *promoter*MAL2, isolate #1 | This work |
| JKC1780 | JKC1772 | *pho84::ARG4/pho84::HIS1 SOD3/FRT-promoterMAL2-SOD3* | JKC1772 after recombination of FRTs following FLP induction | This work |

**S2 Table.** Plasmids used in this Study

| Plasmid | Description | Source (Reference) |
| --- | --- | --- |
| *pGFP-HIS1* |  | (4) |
| pJK1027 | pAU34 with *URA3* disrupted by *Ag<sub>promoter</sub>TEF1-NAT1-Ag<sub>terminator</sub>TEF1* | (5) |
| pJK1085 | *CaTPK1* ligated into *XmaI* sites of pJK1027, used as cloning vector to replace *TPK1* with GFP. | This work |
| pJK1323 | *pACT1-GFP*, product of fjk1615/rjk1633 using *pGFP-HIS1* as template, ligated into *XmaI*/*Clal* sites of pJK1085. | This work |
| pJK1000 | *FLP-NAT1-tetO-PES1*, derived from Litmus 28 (NEB). Used as a template for sub-cloning *SOD3* homology sequences to generate *tetO-SOD3*. | (2) |
| pJK1351 | *FLP-NAT1-tetO-SOD3*, derived from pJK1000. Product of fjk1821 and rjk1822 using SC5314 genomic DNA as template was ligated into *SacII*/*NcoI* sites of pJK1000; product of fjk1819 and rjk1820 using SC5314 genomic DNA as template was ligated into *KpnI*/*Apal* sites of pJK1000. | This work |
| pJK896 | *FLP-NAT1-pMAL2-PES1* | (2) |
| pJK1353 | *FLP-NAT1-pMAL2-SOD3*, derived from pJK1351, *MAL2* promoter was excised from pJK896 using *NotI*/*SacII*, and ligated into p1351 that was digested at the same sites. | This work |

**S3 Table.**
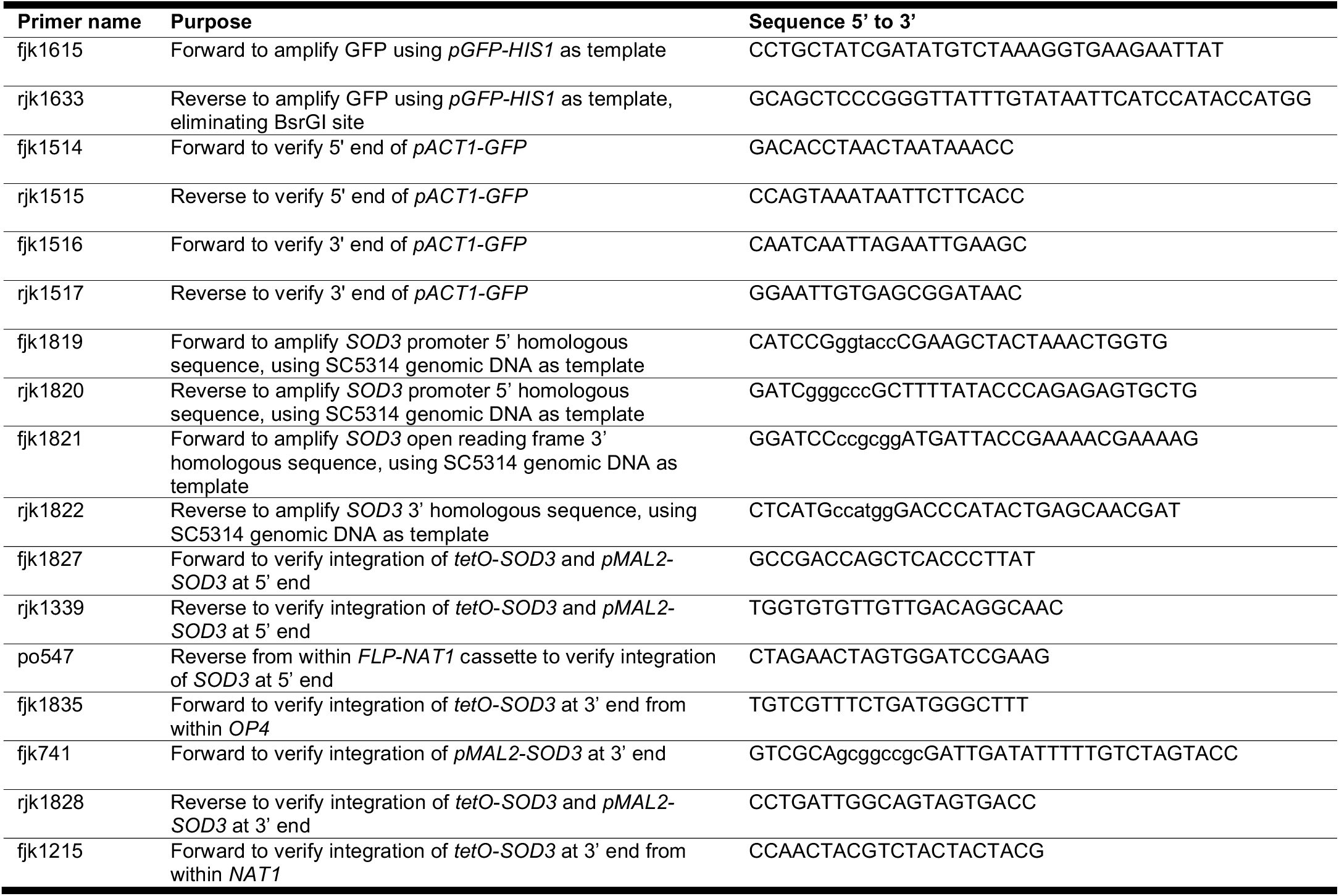
Primers used in this study

**Table S4.** Antibodies used in this study

| Purpose | Antigen recognized | Species | Source or Reference |
| --- | --- | --- | --- |
|  | Hog1 | mouse | Santa Cruz Biotechnology, cat. # sc-165978 |
|  | Phosphorylated Hog1 | rabbit | Cell Signaling Technology, cat. #4511 |
|  | Sod3 | rabbit | Valeria C. Culotta |
| loading control | PSTAIRE | rabbit | Santa Cruz Biotechnology, cat. # sc-53 |
| loading control | Tubulin | rat | Abcam, cat. # ab6161 |
| loading control | Histone H3 | rabbit | Cell Signaling Technology, cat. #4499 |
| secondary | Mouse Ig |  | Santa Cruz Biotechnology, cat. #sc-516102 |
| secondary | Rabbit Ig | bovine | Cell Signaling Technology, cat. # 7074s |
| secondary | Rat Ig | goat | Santa Cruz Biotechnology, cat. #97057 |
| secondary | Goat Ig | donkey | Santa Cruz Biotechnology, cat. #sc-2020 |

